# A genetic code change in progress: tRNA-Leu(CAG) is conserved in most *Saccharomycopsis* yeast species but is non-essential and does not compete with tRNA-Ser(CAG) in translation

**DOI:** 10.1101/2023.07.05.547863

**Authors:** Eoin Ó Cinnéide, Caitriona Scaife, Eugene Dillon, Kenneth H. Wolfe

## Abstract

In the yeast genera *Saccharomycopsis* and *Ascoidea*, nuclear genes use a non-standard genetic code in which CUG codons are translated as serine instead of leucine, due to a tRNA-Ser with the unusual anticodon CAG. However, some species in this ‘CUG-Ser2’ clade also contain an ancestral tRNA-Leu gene with the same anticodon. One of these species, *Ascoidea asiatica*, has been shown to have a stochastic proteome in which proteins contain approximately 50% Ser and 50% Leu at CUG codon sites, whereas previously examined *Saccharomycopsis* species translate CUG only as Ser. Here, we investigated the presence, conservation, and possible functionality of the tRNA-Leu(CAG) gene in the genus *Saccharomycopsis*. We analyzed the genomes of 33 strains, including almost all known species of *Saccharomycopsis*, and found that most of them contain both tRNA-Ser(CAG) and tRNA-Leu(CAG) genes. The tRNA-Leu(CAG) gene is evolving faster than tRNA-Ser(CAG) and it has been lost in two species, *S. microspora* and *S. synnaedendra*. We deleted the single tRNA-Leu(CAG) gene in *S. capsularis* and found that it is not essential. Bioinformatic analysis suggested that some CUG codon sites in *Saccharomycopsis* species may be translated as Leu, specifically in genes with functions in meiosis or sporulation, but mass spectrometry of sporulating *S. capsularis* and *S. fermentans* cultures showed only CUG-Ser translation. Cloverleaf structures of tRNA-Leu(CAG) from all *Saccharomycopsis* species contain mutations that are likely to make them non-functional in translation, but the evolutionary conservation of the gene leads us to propose that it has been retained for an unknown non-translational role.

## Introduction

The genetic code was initially thought to be an immutable frozen accident (Crick 1968; Osawa 1995), because reassignment of a codon’s meaning from one amino acid to another, or to a stop codon, would have a wide-ranging effect similar to mutating every gene in which the codon occurs. Despite this, it is now well established that several codon reassignments have occurred during evolution (Keeling 2016), especially in mitochondrial genomes. Some UGA or UAG codons, which are normally stop codons, are translated as a 21st amino acid selenocysteine or pyrrolysine respectively in certain genes, especially in some methanogenic archaeal species (Rother and Krzycki 2010; Yuan et al. 2010; Prat et al. 2012). Evolutionary reassignments of a codon from one amino acid to another (‘sense-to-sense’ reassignments) are much rarer than reassignments of stop codons to sense codons. In bacteria, a recent extensive computational screen identified six probable instances of sense-to-sense reassignment among 250,000 bacterial genomes examined, all of which involved reassignment of arginine codons (CGG, CGA or AGG) to other amino acids (Met, Gln or Trp) (Shulgina and Eddy 2021).

Among all eukaryotes, only three cases of sense-to-sense codon reassignments have been discovered. They all occurred in budding yeasts (subphylum Saccharomycotina), and they all involved reassignment of the codon CUG, which is translated as leucine in the standard genetic code. Two separate yeast clades, called the CUG-Ser1 and CUG-Ser2 clades (Krassowski et al. 2018), independently reassigned the codon CUG from Leu to Ser. The CUG-Ser1 clade is large (families Debaryomycetaceae and Metschnikowiaceae) and includes the pathogen *Candida albicans* (Kawaguchi et al. 1989; Santos and Tuite 1995; Sugita and Nakase 1999). The CUG-Ser2 clade is much smaller and includes only two genera: *Ascoidea* and *Saccharomycopsis* (Krassowski et al. 2018; Muhlhausen et al. 2018; Junker et al. 2019). A small third yeast clade, which contains three genera including the species *Pachysolen tannophilus*, reassigned CUG from Leu to Ala (Mühlhausen et al. 2016; Riley et al. 2016; Krassowski et al. 2018; Muhlhausen et al. 2018). Each of these reassignments was mediated by duplication of an ancestral tRNA-Ser or tRNA-Ala gene, followed by mutation of the anticodon to become CAG, allowing translation of CUG codons by a tRNA that is charged with Ser or Ala. The CUG-Ser1 and CUG-Ser2 clades arose by duplication and anticodon mutation of two different tRNA-Ser genes, one from the four-codon box (UCN) and one from the two-codon box (AGY) of serine codons (Krassowski et al. 2018). Interestingly, serine and alanine aminoacyl synthetases (SerRS and AlaRS) are the only aminoacyl synthetases that do not use bases in the anticodon to recognize a tRNA’s identity when charging it (Giegé et al. 1998), so Ser and Ala tRNAs can remain functional with any anticodon sequence, whereas tRNAs for the other 18 amino acids cannot. For this reason, it is probably easier to reassign sense codons to Ser and Ala than to any other amino acids (Kollmar and Mühlhausen 2017a, b).

Of the three yeast clades with genetic code reassignments, the CUG-Ser2 clade (*Saccharomycopsis* and *Ascoidea*) is distinct because it is the only clade in which some species appear to contain genes for both the novel tRNA (tRNA-Ser(CAG)) and the original tRNA (tRNA-Leu(CAG)) (Krassowski et al. 2018; Muhlhausen et al. 2018; Junker et al. 2019). In the two other clades, tRNA-Leu(CAG) has been lost. The presence of both genes in the same genome may indicate that the evolutionary process of changing the genetic code in the CUG-Ser2 clade is still in progress, whereas it has finished in the other two clades. However, the functional role of tRNA-Leu(CAG) is not very clear. In *Ascoidea asiatica*, genes for both tRNA-Ser(CAG) and tRNA-Leu(CAG) are present in the genome and this species has been found to have a stochastic proteome in which CUG codons in mRNAs are translated randomly as either Ser or Leu, so both of the tRNAs are used for translation (Muhlhausen et al. 2018). In contrast, although both of the tRNA genes are also present in at least four *Saccharomycopsis* species (*S. capsularis*, *S. malanga*, *S. fibuligera*, and *S. schoenii*), proteomic investigation of these species by mass spectrometry indicated that they translate CUG only as Ser (Krassowski et al. 2018; Muhlhausen et al. 2018; Junker et al. 2019). This observation made it doubtful that tRNA-Leu(CAG) has a role in translation in *Saccharomycopsis* species, but Shulgina and Eddy (2021) recently showed that in *S. malanga* both tRNA-Leu(CAG) and tRNA-Ser(CAG) are transcribed and aminoacylated, which suggests that they can both be used in translation.

In this study, our goal was to investigate the presence, conservation, and possible function of tRNA-Leu(CAG) genes in *Saccharomycopsis* species. We sequenced the genomes of strains spanning the whole genus, constructed a phylogenomic tree, and compared their tRNA gene content. We developed a CRISPR/Cas9 gene editing technique for one species, *S. capsularis*, and deleted its tRNA-Leu(CAG) gene to test its essentiality. We also designed a bioinformatic screen of sequence conservation in *Saccharomycopsis* genes to search for genes that contain CUG codons at sites of conserved Leu residues, which potentially could require CUG-Leu translation. This screen led us to hypothesize that CUG-Leu translation might occur during sporulation in *Saccharomycopsis*, but this hypothesis was not supported by our mass spectrometry experiments. Based on our analyses, we conclude that the tRNA-Leu(CAG) gene in *Saccharomycopsis* is not used for translation but it may still retain a role in an unidentified non-translational process.

## Results

### Phylogenomic tree of the genus *Saccharomycopsis*

We analyzed genome sequence data from 20 species (33 strains) of yeasts in the genera *Saccharomycopsis* and *Ascoidea*, comprising 23 strains that were newly sequenced for this study using Illumina short-read sequencing, and 10 strains whose genome sequences were downloaded from NCBI (Table S1). The dataset includes almost all the known species in both genera. The average genome assembly size is 16.3 Mb, with a range from 12.2 Mb to 22.3 Mb (Table S1). We did not include known hybrid strains with double-size genomes, such as *S. fibuligera* strain KJJ81 (Choo et al. 2016), in the analysis.

To investigate the phylogenetic relationships among the 33 strains, we identified a set of 1,227 single-copy orthologous protein groups by using OrthoFinder (Emms and Kelly 2019). The proteins in each group were aligned, trimmed, and concatenated, yielding a supermatrix with 567,098 sites. IQ-TREE (Minh et al. 2020) found the best Maximum Likelihood model to be LG+F+R5, and almost all branches were completely supported by ultrafast bootstrapping and the Shimodaira-Hasegawa Approximate Likelihood Ratio Test (SH-ALRT) (Shimodaira 2002).

Our phylogenomic tree is shown in Figure 1A. The tree was rooted by assuming that *Saccharomycopsis* is monophyletic and *Ascoidea* is an outgroup to it. The OrthoFinder analysis, based on gene duplication events, also supported this position for the root. The tree confirms several clusters of *Saccharomycopsis* species that were also apparent in a previous tree published by Jacques et al. (2014) based on sequencing a small number of loci (D1/D2 26S rDNA region, SSU rDNA, and the gene *TEF1*; Figure S1 of (Jacques et al. 2014)). Both analyses have a large cluster consisting of *S. schoenii*, *S. oosterbeekiorum*, *S. javanensis*, *S. fermentans*, and *S. babjevae*, and the following monophyletic pairs of species: *S. crataegensis* with *S. amapae*; *S. synnaedendra* with *S. microspora*; and *S. guyanensis* with *S. fodiens*. However, the positions of these groups relative to each other is substantially different between our tree and that of Jacques et al. (2014). Although many of the internal branches in our phylogenomic tree are short (Fig. 1A), they have strong statistical support whereas many internal branches in the rDNA/*TEF1* tree had poor support (Jacques et al. 2014). Our phylogenomic dataset is much larger than the rDNA/*TEF1* dataset and is therefore expected to give a more reliable tree. It indicates that the deepest-branching species in the genus is *S. selenospora*. It also places the economically important species *S. fibuligera* (used to manufacture the alcoholic drink nuruk) as a sister clade to *S. capsularis* and *S. malanga* (Fig. 1A). When compared to another recent phylogenomic tree constructed from a smaller number of *Saccharomycopsis* genome sequences (Yuan et al. 2021), our tree differs in the location of the root and in the position of *S. fodiens* and the unnamed *Saccharomycopsis* species UWOPS 91-127.1 relative to the other clades, but again it has much stronger statistical support.

**Figure 1.**
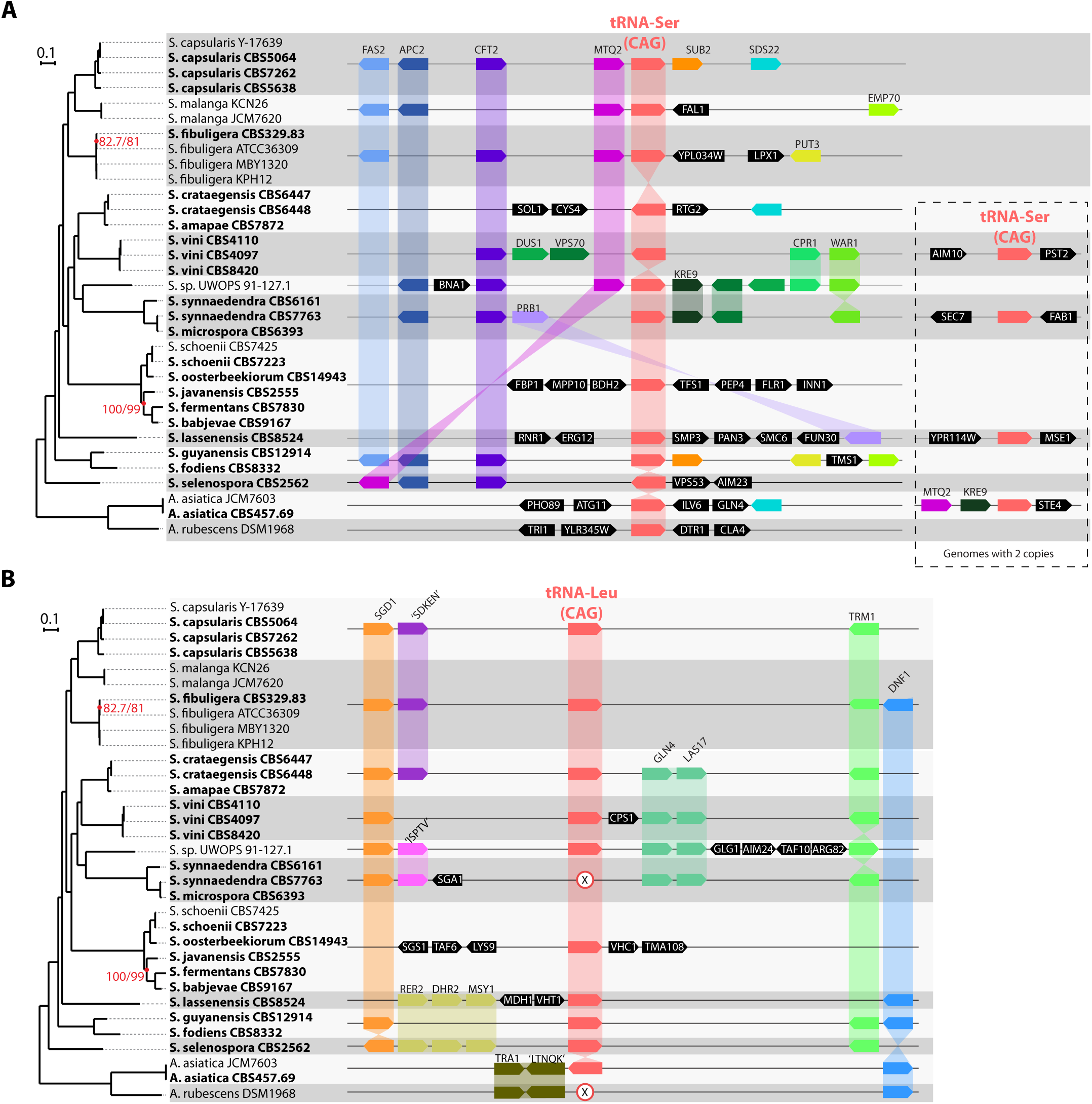
Phylogenomic tree of *Saccharomycopsis* and *Ascoidea* species, and synteny relationships around (A) the tRNA-Ser(CAG) genes, and (B) the tRNA-Leu(CAG) genes. The same tree is shown in both panels. Strain names are indicated beside species names, and genomes sequenced in this study are indicated in bold. All branches except two were supported by 100% ultra-fast bootstrapping, and 100% SH-alrt; the two branches not fully supported are indicated by the red text. (A) Synteny relationships around tRNA-Ser(CAG) loci. Gray and white horizontal bands indicate groups of strains or species with identical gene arrangements, and only one gene diagram is shown for each group. Some genomes contain a second tRNA-Ser(CAG) gene, shown in the dashed box. In *A. asiatica*, one sequenced strain (JCM7603) contains two tRNA-Ser(CAG) genes but the other (CBS457.69) contains only one. The copy that is only in JCM7603 shares neighbors (*MTQ2* and *KRE9*) with *Saccharomycopsis* species. (B) Synteny relationships around the tRNA-Leu(CAG) locus. X symbols mark the absence of a tRNA-Leu(CAG) gene in genomes that have lost this gene. Genes are named according to their *S. cerevisiae* orthologs where possible. Other genes are named by conserved sequence motifs they contain.

Our phylogenetic analysis also detected one strain that appears to have been misidentified. CBS 7763, which was deposited in culture collections as *S. synnaedendra*, is very closely related to the type strain of *S. microspora* (CBS 6393) and groups with it rather than with the type strain of *S. synnaedendra* (CBS 6161) (Fig. 1A), so CBS 7763 therefore appears to be a strain of *S. microspora*. This misidentification was also previously reported by Quintilla et al. (2018) using MALDI-TOF mass spectrometry as a tool for species identification.

### The tRNA-Leu(CAG) gene is present in most of the Ser2 clade and is at an ancestral location

We used tRNAscan-SE (Lowe and Eddy 1997) and BLASTN searches with manual annotation to identify tRNA genes in each genome sequence, and then examined the local gene order around these loci. Most of the species contain both a tRNA-Ser(CAG) gene (Fig. 1A) and a tRNA-Leu(CAG) gene (Fig 1B).

All the species and strains examined contain at least one gene for tRNA-Ser(CAG), the novel tRNA that allows translation of CUG codons as Ser (Fig. 1A, right panel). Synteny of the nearby protein-coding genes is partially conserved, notably the genes *CFT2* and *APC2* which are neighbors of tRNA-Ser(CAG) in several *Saccharomycopsis* clades. There is a second tRNA-Ser(CAG) gene in four clades, but these genes are at different genomic locations that are not conserved so they appear to be the result of independent gene duplications (Fig. 1A, dashed box).

The gene for tRNA-Leu(CAG), the ancestral tRNA that can potentially translate CUG codons as Leu, is present in 17 of the 20 species so it is broadly conserved (Fig. 1B). It is missing in three species: in *A. rubescens* as previously reported (Krassowski et al. 2018; Muhlhausen et al. 2018), and separately in the species pair *S. synnaedendra* and *S. microspora* (Fig. 1B). Synteny of other genes around the tRNA-Leu(CAG) locus is reasonably well conserved, except in the clade containing *S. fermentans* where rearrangements have occurred. The conserved synteny indicates that the tRNA-Leu(CAG) genes in each species are orthologs, except possibly in the *S. fermentans* clade. Synteny in this region is also conserved in the three species that have lost the tRNA gene, and we did not find any pseudogene or relic of it. The protein-coding gene *TRM1*, which is beside or close to tRNA-Leu(CAG) in most *Saccharomycopsis* species (Fig. 1B), is also located beside tRNA-Leu(CAG) in yeasts in the genera *Wickerhamomyces* and *Cyberlindnera* (family Phaffomycetaceae) (Krassowski et al. 2018). Those genera are in the CUG-Leu1 clade and translate CUG as Leu, so the shared adjacency of tRNA-Leu(CAG) to *TRM1* suggests that this is the ancestral location of the tRNA gene, i.e. that the tRNA-Leu(CAG) gene in most *Saccharomycopsis* species is a surviving ortholog of the functional tRNA-Leu(CAG) gene of *Wickerhamomyces* and *Cyberlindnera*.

### The tRNA-Leu(CAG) gene is non-essential in *S. capsularis*

To test whether the tRNA-Leu(CAG) gene is essential in a *Saccharomycopsis* species, we deleted it by a CRISPR-Cas9 approach. We used *S. capsularis* strain NRRL Y-17639 for this experiment because it is the type strain of the entire genus. No plasmid-based CRISPR system has been developed for any *Saccharomycopsis* species, so instead we used an approach similar to Grahl et al. (2017), in which ribonucleoproteins (RNPs) containing Cas9 protein and a single guide RNA (sgRNA) were co-transformed into competent cells together with a repair template (double-stranded DNA). We used the Alt-R CRISPR-Cas9 system from Integrated DNA Technologies. We first tested the system by disrupting the *ADE2* gene of *S. capsularis* NRRL Y-17639 with either a kanamycin, nourseothricin, or hygromycin drug resistance cassette, each of which had been engineered to lack CUG codons. We recovered *ade2* mutant colonies at high efficiency, using repair templates designed to disrupt *ADE2* (Fig. S1).

We then used the same approach to delete the tRNA-Leu(CAG) gene, including approximately 200 bp upstream and downstream, in the wildtype NRRL Y-17639 genetic background (Fig. 2A). Kanamycin-resistant transformants, from two independent experiments, were verified by PCR (Fig. 2B) and Sanger sequencing of the entire recombinant locus. Nine tRNA-Leu(CAG) deletion mutants were obtained. Transformants were viable and there were no obvious morphological differences (of colonies, or of cells examined by light microscopy) between the deletion strains and the wildtype strain, on YPD or synthetic complete (SC) media. In liquid growth assays, there was no significant difference between deletion and wildtype strains at either 25°C or 37°C, which are respectively the optimal and maximal growth temperature for *S. capsularis* (Kurtzman et al. 2011) (Fig. 2C).

**Figure 2.**
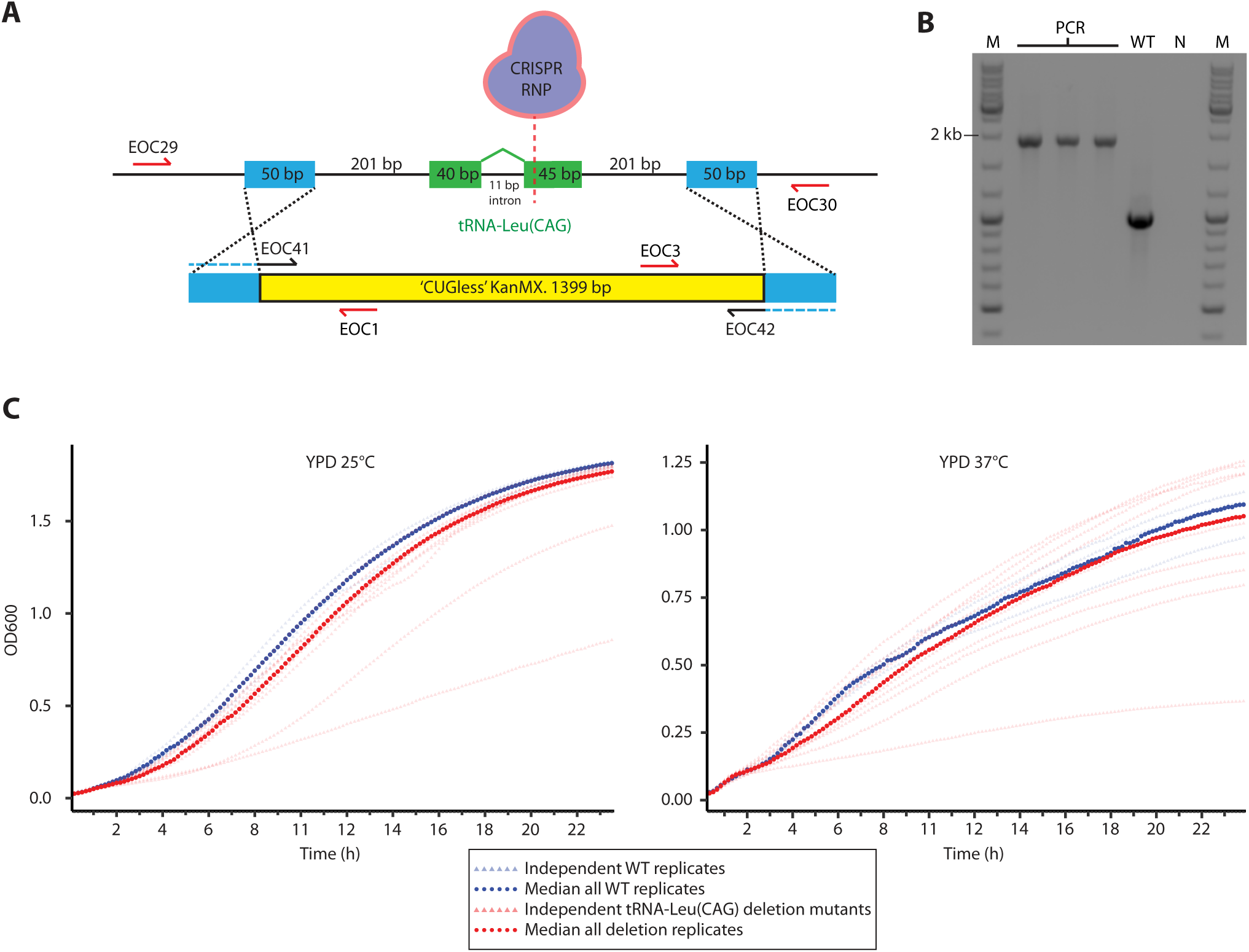
Deletion of tRNA-Leu(CAG) in *S. capsularis* is viable. (A) Scheme for deletion of the tRNA-Leu(CAG) gene of *S. capsularis* NRRL Y-17639 by CRISPR-Cas9 RNPs. The repair template containing a kanamycin resistance cassette (KanMX, modified to lack CUG codons) was amplified from a plasmid using primers with 50-nucleotide tails (blue) to generate homology arms matching the regions upstream and downstream of the tRNA-Leu(CAG) gene. CRISPR RNPs containing a guide RNA targeting exon 2 of the tRNA-Leu(CAG) gene were assembled *in vitro* and electroporated into competent cells. Transformants were screened by junction PCR using primer pairs EOC29 + EOC1 and EOC3 + EOC30. (B) PCR confirmation of three tRNA-Leu(CAG) deletion strains, by amplification of the entire recombinant locus with primer pair EOC29 + EOC30. The 3 lanes marked PCR are transformant colonies with the deletion; WT is a wildtype NRRL Y-17639 control; N is a negative control; M is a size marker ladder. (C) Growth curves comparing the nine independent tRNA-Leu(CAG) deletion strains (red), to three biological replicates of the wildtype strain (blue), at 25°C and 37°C. Median values at each timepoint are shown in the darker shade.

*S. capsularis* strain NRRL Y-17639 does not sporulate (see below), so we also attempted to delete the tRNA-Leu(CAG) gene in three other strains of *S. capsularis* that sporulate well (CBS5064, CBS5638 and CBS7262). However, we were unsuccessful in each case, even with the length of homology arms increased to 500 bp or 1 kb. In these strains, drug-resistant transformants were obtained but we never detected successful integration at the target locus by PCR. Thus we conclude that in these other strains, homologous recombination was less efficient than in NRRL Y-17639 and the drug resistance marker integrated at ectopic locations in the genome.

### Bioinformatic inference of potential CUG-Leu sites in genes

The tRNA-Leu(CAG) gene is broadly conserved across the genus *Saccharomycopsis* (Fig. 1B) despite the apparent lack of use of this tRNA to translate CUG codons in several species whose proteomes have been examined by mass spectrometry (Krassowski et al. 2018; Muhlhausen et al. 2018; Junker et al. 2019). We therefore hypothesized that retention of the tRNA-Leu(CAG) gene might be due to a handful of specific CUG codon sites, *i.e.* that a small number of genes might contain a CUG codon at a site where translation as serine leads to a non-functional protein, so these sites need to be translated as Leu and not Ser. Even if only a very small number of such ‘essential CUG-Leu’ sites exists, they would generate selection pressure to retain a functional tRNA-Leu(CAG) in the genome. If a leucine residue at these sites is essential for function of the protein, and the protein is essential for viability, the only way that loss of the tRNA-Leu(CAG) gene from the genome can be tolerated is if the site first mutates to become a different leucine codon (CUU, CUC, CUA, UUA or UUG) instead of CUG. We further hypothesized that essential CUG-Leu sites might be present in only a specific functional subset of genes, meaning that expression of tRNA-Leu(CAG) might be restricted to a specific growth condition or stage of the cell cycle, explaining why its use has not been detected by proteomics.

To search computationally for possible essential CUG-Leu sites in genes, we constructed multiple amino acid sequence alignments (MSAs) of proteins in 4,187 orthogroups (sets of orthologous proteins or genes) from *Saccharomycopsis* and *Ascoidea*, and analyzed them using a heuristic set of rules (Fig. 3A; see Methods). This heuristic approach leverages the three species that do not have a tRNA-Leu(CAG) gene (*S. synnaedendra*, *S. microspora* and *A. rubescens*). The aim was to find columns in the MSAs that ancestrally may have been a CUG codon, and that have been conserved as a Leu residue in the three species unable to translate CUG as Leu anymore. We searched for columns in the MSAs that are Leu in these three species, and that contain a CUG codon in at least one of the other species that have both tRNA genes (Fig. 3A). We found 602 MSAs that have at least one column of this type. For each of these columns we calculated a ‘Column Score’, which is the proportion of CUG codons and non-CUG Leu codons among all the sequences in the alignment and is an approximate measure of the importance of having Leu at this position in the protein. Finally, for any MSAs that contain multiple columns of this type, we summed the Column Scores of each column to generate a ‘CUG-Leu Conservation Score’ for the whole protein represented by that MSA (Fig. 3A), which is an approximate measure of the importance of translating all the CUG codons in this orthogroup’s gene as Leu. Most of the orthogroups have a CUG-Leu Conservation Score < 1, which means that there is less than one well-conserved column of this type in its MSA, whereas a few proteins had scores > 10 (Fig. 3B). There is no correlation between the score and the length of the MSA (Fig. 3B).

**Figure 3.**
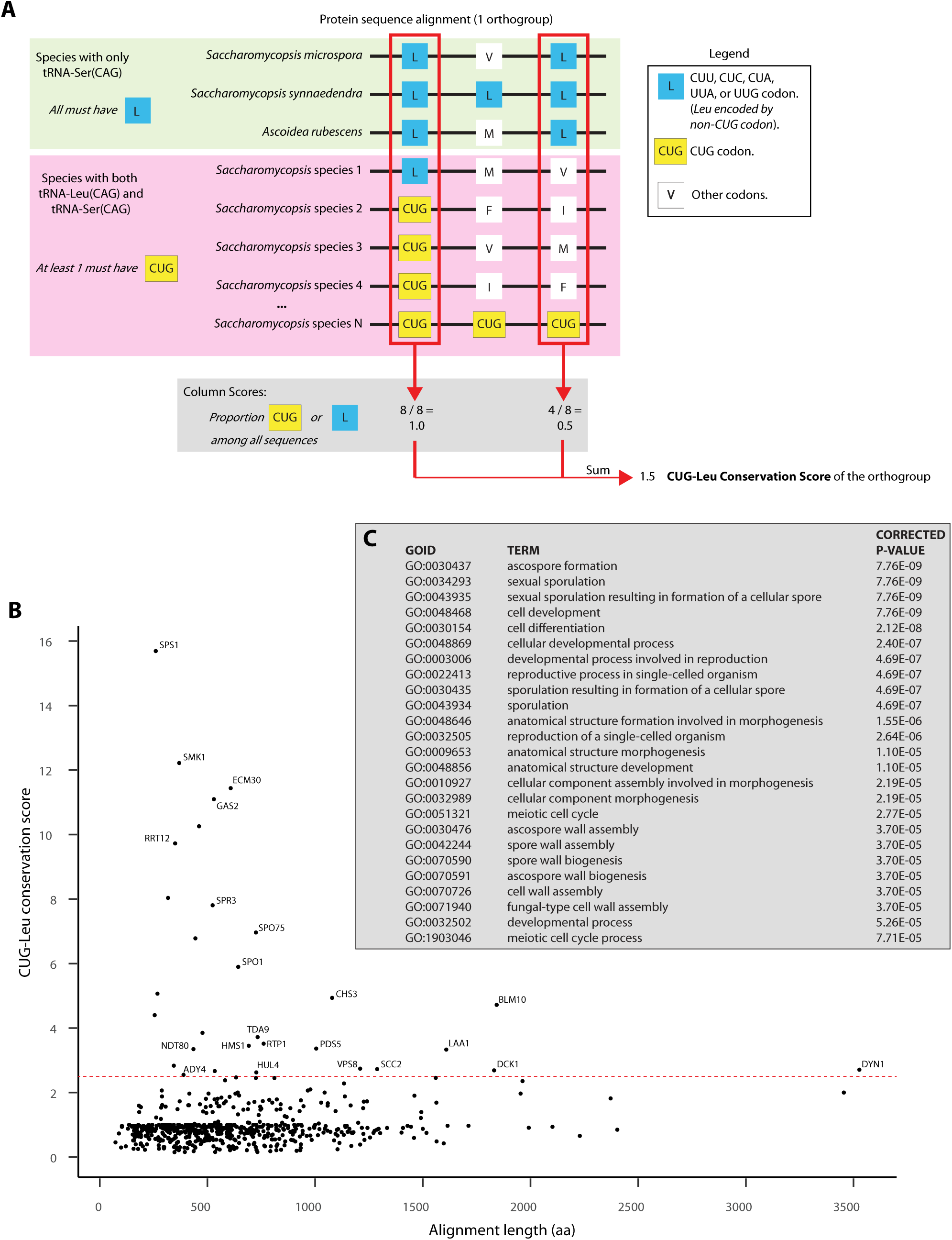
Bioinformatic search for genes with potentially conserved CUG-Leu translation. (A) Summary of the method. The diagram represents a multiple sequence alignment (MSA) of proteins in an orthogroup from *Saccharomycopsis* and *Ascoidea* species, with three columns (aligned amino acid sites) shown. The green background indicates the three species that have no tRNA-Leu(CAG) gene, and the pink background indicates the larger group of other species that have both tRNAs (up to 30 sequences were present in this group). Red boxes show the two columns that are of interest, because they contain a conserved Leu in the green group, and they contain at least one CUG codon in the pink group. The gray box shows how a Column Score is calculated for each column of interest, and then the scores from all columns of interest are summed to generate the CUG-Leu Conservation Score for the orthogroup. (B) Scatterplot of CUG-Leu Conservation Scores versus MSA length, for the 602 orthogroups that contain ≥25 sequences and at least one column of interest. The dashed red horizontal line shows the top 5% of scores, and orthogroups above this line are named according to their *S. cerevisiae* ortholog (assigned by a BLASTP reciprocal best hit method; absence of a name indicates that no ortholog was found). (C) Gene Ontology terms enriched in genes with the highest 5% of CUG-Leu Conservation Scores, compared to background (a set of 519 *S. cerevisiae* genes, which are the identifiable orthologs of the 602 orthogroups plotted in B). GO term enrichment was calculated using the SGD GO Term Finder, and all terms with corrected P-values < 10^-4^ are shown.

We identified *S. cerevisiae* orthologs for as many of the 602 orthogroups as possible, using a reciprocal BLASTP approach, and then investigated whether particular Gene Ontology (GO) categories were over-represented among the proteins with high CUG-Leu Conservation Scores, using the GO annotations of their *S. cerevisiae* orthologs and the GO Term Finder tool on the *Saccharomyces* Genome Database website (Boyle et al. 2004; Engel et al. 2022). This analysis showed that many GO categories associated with yeast sporulation and meiosis are enriched, with very high statistical significance, among the genes with the highest 5% of CUG-Leu Conservation Scores (Fig. 3C). Strikingly, almost every orthogroup with a high CUG-Leu Conservation Score (Fig. 3B) has a role in sporulation. They include *SPS1* (kinase required for prospore membrane closure), *SMK1* (kinase required for prospore membrane development), *GAS2* (involved in spore wall assembly), *RRT12* (formation of dityrosine layer of spore wall), *SPR3* (septin involved in sporulation), *SPO75* (required for spore wall formation), *SPO1* (required for prospore membrane assembly), and *CHS3* (required for spore wall chitosan synthesis) (Engel et al. 2022). Each of these eight sporulation genes has a CUG-Leu Conservation Score >4, whereas there are only two identified genes (*ECM30* and *BLM10*) with scores >4 that are not involved in sporulation (Fig. 3B). The statistical enrichment of GO terms related to sporulation, meiosis, or development was also confirmed using two other bioinformatics tools: YeastEnrichr (Kuleshov et al. 2019; Xie et al. 2021) and Gene Set Enrichment Analysis (GSEA; (Subramanian et al. 2005)) (Fig. S2).

### Tandem mass spectrometry of sporulating cultures

Sporulation is a specialized part of the yeast life cycle in which diploid cells undergo meiosis and produce four haploid spores, and in most yeast species including *Saccharomycopsis* it occurs when induced by conditions such as nutrient starvation (Kreger-van Rij 1964; Neiman 2011). Many genes required for sporulation are not expressed during normal vegetative (mitotic) growth (Chu et al. 1998; Brar et al. 2012). Because of the enrichment for sporulation functions in genes with high CUG-Leu Conservation Scores (Fig. 3), we hypothesised that perhaps tRNA-Leu(CAG) is required for translation during sporulation or meiosis. To investigate this possibility, we performed tandem mass spectrometry (LC-MS/MS) on yeast cultures grown in vegetative or sporulation conditions, to determine how they translate CUG codons.

*S. capsularis* strain NRRL Y-17639 (also called CBS 2519) is the only genetic background in which we were able to delete the tRNA-Leu(CAG) gene. Unfortunately, although this strain has previously been reported to sporulate (Lodder and Kreger-van Rij 1952; Kurtzman and Smith 2011), we were unable to induce it to sporulate in any conditions that we tried, so we cannot directly test whether tRNA-Leu(CAG) is essential for sporulation. Instead, we took a proteomics approach to investigate whether translation of CUG codons as leucine could be detected in sporulating cultures of *S. capsularis*, using three other wildtype strains that sporulate well (CBS5064, CBS5638 and CBS7262). In addition, we compared the proteomes of wildtype NRRL Y-17639 and its tRNA-Leu(CAG) deletion mutant, grown in conditions that induce sporulation in the other strains. We also analyzed the proteome of a second species, *S. fermentans* CBS7830, chosen because its tRNA-Leu(CAG) gene does not share synteny with the *S. capsularis* gene (Fig. 1B).

We grew each strain on solid media, in either sporulation media (YM agar, 25°C, 24 days for all *S. capsularis* strains; 5% ME agar, 25°C, 25 days for *S. fermentans*) or vegetative media (YPD agar, 25°C, 3 days). Cells were scraped from the plates, resuspended in sterile water, and light microscopy was used to verify sporulation. For *S. capsularis* the cell samples were lysed and analyzed by LC-MS/MS on a Q Exactive Quadrupole Orbitrap mass spectrometer (Thermo Fisher). An average of 13,800 peptides were sequenced per strain, resulting in sampling of an average of 296 CUG codon sites in genes per strain. The only significant signal was of CUG translation as Ser (Fig. 4A-E). There was no difference in translation patterns between sporulation and vegetative conditions, and no difference among the strains including the tRNA-Leu(CAG) knockout strain. Translation of CUG as Leu was not detected above the level of background seen for other amino acids.

**Figure 4.**
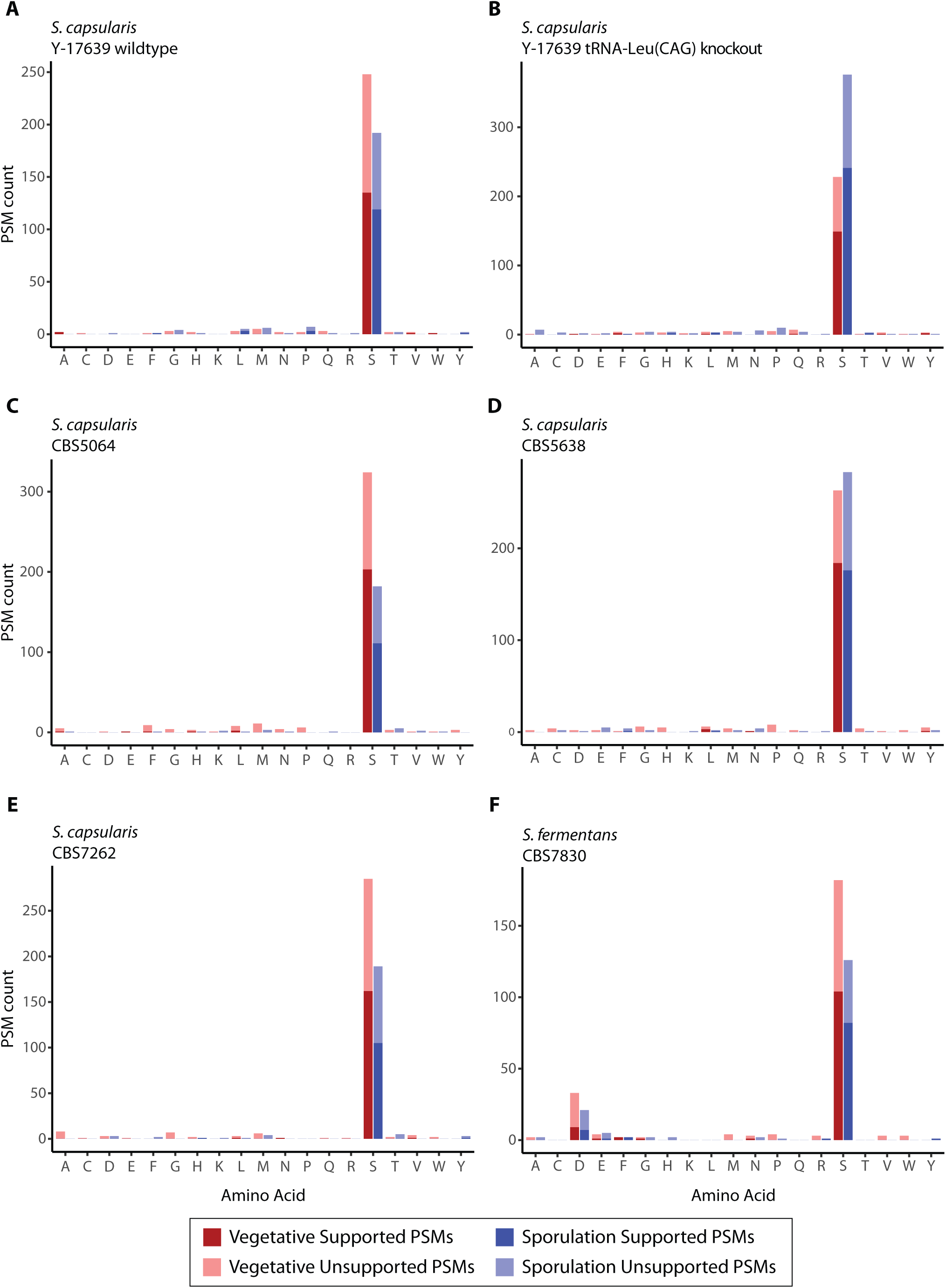
LC-MS/MS analysis of CUG codon translation in *Saccharomycopsis* strains grown in sporulation and vegetative growth conditions (see Methods). Number of peptide spectral matches (PSMs) spanning CUG codons are plotted, for each possible amino acid translation, for six different *Saccharomycopsis* strains: (A) *S. capsularis* NRRL Y-17639 wildtype; (B) *S. capsularis* NRRL Y-17639 tRNA-Leu(CAG) knockout; (C) *S. capsularis* CBS5064; (D) *S. capsularis* CBS5638; (E) *S. capsularis* CBS7262; (F) *S. fermentans* CBS7830. Supported PSMs have either fragment ion b_n_ or y_n_ at the CUG codon (position n) in the peptide, as well as the respective preceding ion in the series (b_n-1_ or y_n-1_). Unsupported PSMs include spectra that match predicted peptides but do not have b- or y- ion support for the particular CUG codon in question.

For *S. fermentans*, we performed SDS-PAGE gel fractionation of total lysed proteins. Each digested gel fraction was run on a Bruker timsTOF instrument, and an average of 8,600 peptides were sequenced and 205 CUG codon sites were sequenced per sample. Again, in both conditions, most CUG codons were translated as Ser and we did not see any well-supported CUG-Leu translation. Surprisingly, approximately 13% of CUG codons in *S. fermentans* appeared to be translated as aspartic acid (D), in both sporulation and vegetative conditions (Fig. 4F). However, we suspect that this result is an artifact caused by misidentification of formylated Ser residues as Asp residues, as described previously (Mordret et al. 2019), because their molecular masses are almost identical (115.026945 and 115.026950 respectively, i.e. a difference of 5 × 10^-6^ mass units). This artifact appeared in the Bruker timsTOF data but not in the data acquired on the Thermo Q-Exactive used for the *S. capsularis* experiments which has a higher resolving power.

### tRNA-Leu(CAG) has structural irregularities in all *Saccharomycopsis* species

The finding that *Saccharomycopsis* species appear to translate CUG codons only as serine, even during sporulation, prompted us to examine the tRNA-Leu(CAG) and tRNA-Ser(CAG) genes and their predicted tRNA cloverleaf structures in more detail. We found that the leucine tRNA is much more poorly sequence conserved than the serine tRNA, with only 22 completely conserved nucleotide sites as compared to 42 (Fig. S3A). The high diversity of tRNA-Leu(CAG) sequences is also apparent in a phylogenetic tree (Fig. S3B).

Examination of the predicted cloverleaf structures of tRNA-Leu(CAG) genes revealed unusual features in almost every species (Fig. 5), when compared to data compiled in the tRNAviz database of tRNA structures from >1500 species (Lin et al. 2019). Whereas the functional tRNA-Leu(CAG) of *A. asiatica* has a cloverleaf structure typical of a eukaryotic tRNA-Leu and contains the correct nucleotide at every site that is highly conserved among eukaryotic leucine tRNAs (Fig. 5A,B), the structures predicted in all the *Saccharomycopsis* species have irregularities that make us doubt that they are functional tRNAs. First, there is no variable arm in tRNA-Leu(CAG) of *S. vini* (Fig. 5C), whereas an extended variable arm (≥ 5 nt) is a hallmark of tRNA-Leu and tRNA-Ser, but not of other tRNA types. Second, some of the predicted *Saccharomycopsis* tRNA-Leu(CAG) molecules have unusually large D-loops. In *A. asiatica* the D-loop is 8 nt long corresponding to positions 14-21 in the standard tRNA numbering scheme (Fig. 5A,B; it also has a mismatch of G_13_:A_22_ at the end of the D-stem, as is common in Saccharomycotina tRNA-Leu). The *A. asiatica* D-loop is consistent in size and sequence with the majority of other eukaryotic tRNA-Leu sequences (Lin et al. 2019). However, the D-loop has expanded to 11 nt in *S. capsularis* (Fig. 5D) and *S. malanga*, and to 12 nt in *S. selenospora* (Fig. 5E), making it larger than any conventional eukaryotic tRNA D-loop. The *S. capsularis* D-loop is also unusual by lacking a uracil residue at position 20; this position is normally modified to dihydrouracil, giving the D-loop its name. Third, the predicted *Saccharomycopsis* tRNA-Leu(CAG) molecules have mutations at positions that are very highly conserved among eukaryotic tRNA-Leu molecules, as shown by the red arrows in Fig. 5. In fact, inspection of the sequences in all 16 *Saccharomycopsis* species with tRNA-Leu(CAG) candidates shows that they all contain mutations at positions that are almost universally conserved among eukaryotic tRNA-Leu molecules, and most of them contain multiple mutations of this type (Fig. 6). The *Ascoidea asiatica* tRNA-Leu(CAG) is therefore exceptional in retaining these conserved residues, and in retaining translational functionality.

**Figure 5.**
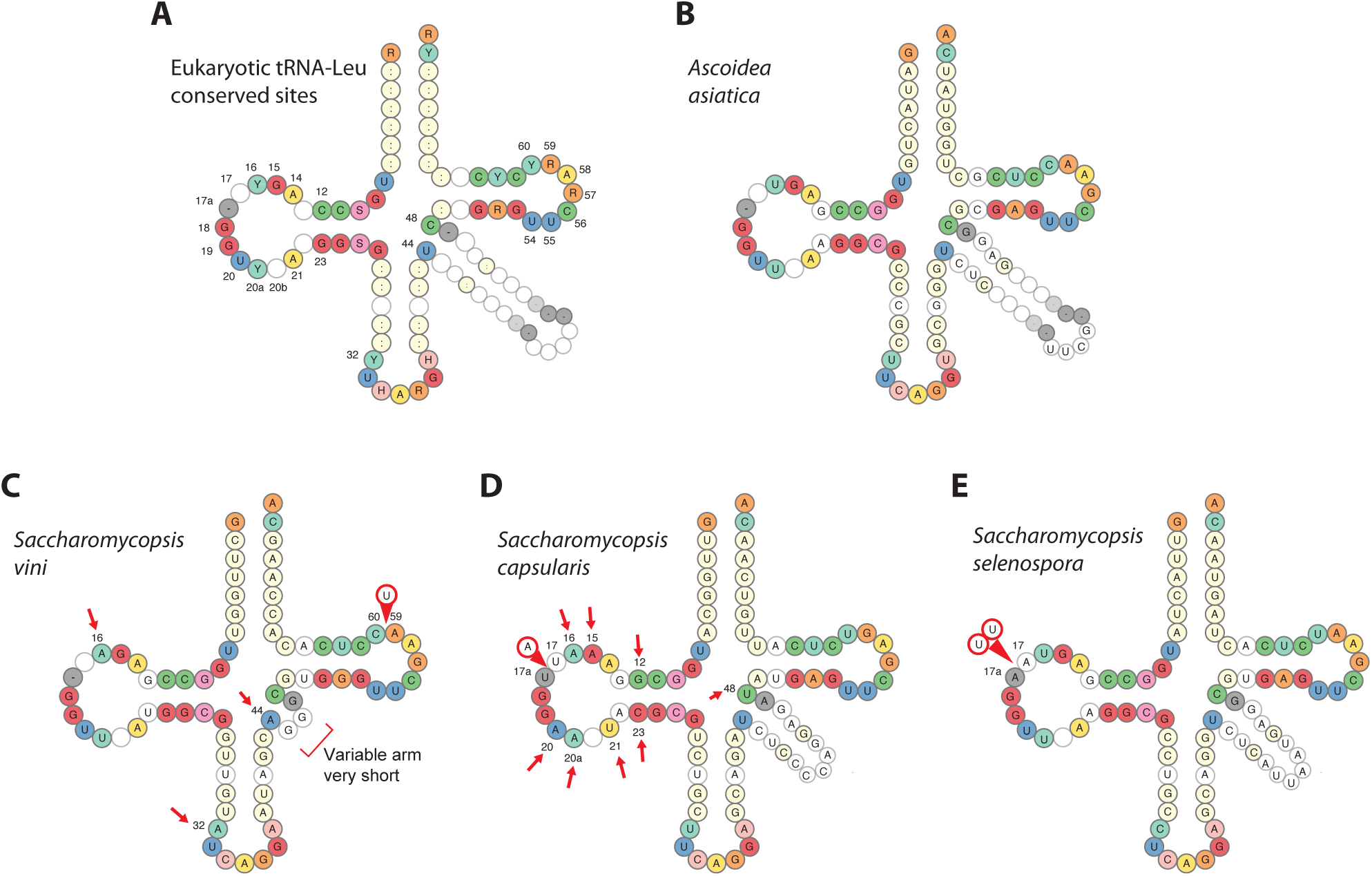
Predicted cloverleaf structures of tRNA-Leu molecules. (A) Nucleotides that are almost universally conserved in eukaryotic tRNA-Leu molecules, from data compiled in the tRNAviz database (Lin et al. 2019). Numbering is shown for some positions, following the standard tRNA numbering scheme (Sekulovski and Trowitzsch 2022). IUPAC ambiguity codes are used. (B-E) Predicted cloverleaf structures of tRNA-Leu(CAG) molecules in *Ascoidea asiatica* and three *Saccharomycopsis* species. Red arrows indicate positions that do not match the highly conserved sites shown in A, and circled bases indicate insertions of bases relative to the standard tRNA structure. *Ascoidea asiatica* is the only CUG-Ser2 clade species that retains all the conserved sites and has no insertions.

**Figure 6.**
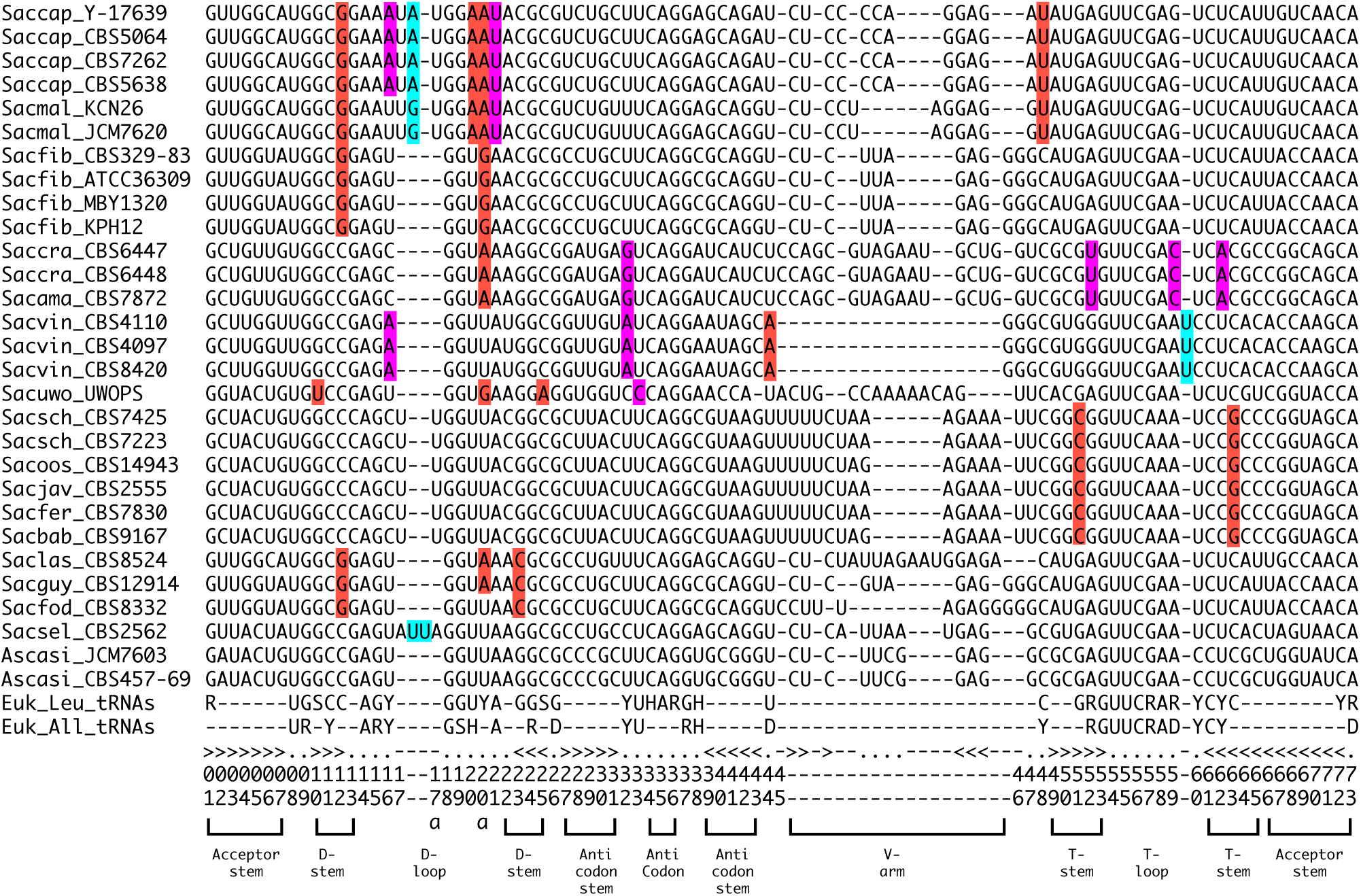
Alignment of tRNA-Leu(CAG) sequences from all sequenced *Saccharomycopsis* and *Ascoidea* species, highlighting variants that may make these genes nonfunctional. Standard tRNA position numbering and secondary structure elements are shown at the bottom. Magenta highlighting shows sites that disagree with nucleotides that are strongly conserved in all eukaryotic tRNAs (Euk_All_tRNAs), and red highlighting shows sites that disagree with nucleotides that are strongly conserved in all eukaryotic leucine tRNAs (Euk_Leu_tRNAs), according to data compiled from thousands of genomes in the tRNAviz database (Lin et al. 2019). Blue highlighting shows insertions of bases in the D-loop and T-loop in some species, making these loops longer than can be accommodated by the standard numbering system.

Despite the apparent defects in the cloverleaf structures, comparative analysis does suggest that the sequences of the tRNA-Leu(CAG) locus have been preserved during evolution to a greater extent than expected by chance. For example, a dot matrix comparison between *S. babjevae* and *S. fermentans* shows that the tRNA locus has been conserved whereas most of the intergenic region that surrounds it (between the protein-coding genes *LYS9* and *VHC1*) has diverged (Fig. 7A). Furthermore, the sequences of the exons of the tRNA gene are almost identical, whereas its intron and the alignable region upstream of exon 1 have diverged (Fig. 7B). Similarly, we previously found that the tRNA-Leu(CAG) gene was better conserved than its flanking regions in a comparison among *S. capsularis*, *S. malanga* and *S. fibuligera* (Krassowski et al. 2018).

**Figure 7.**
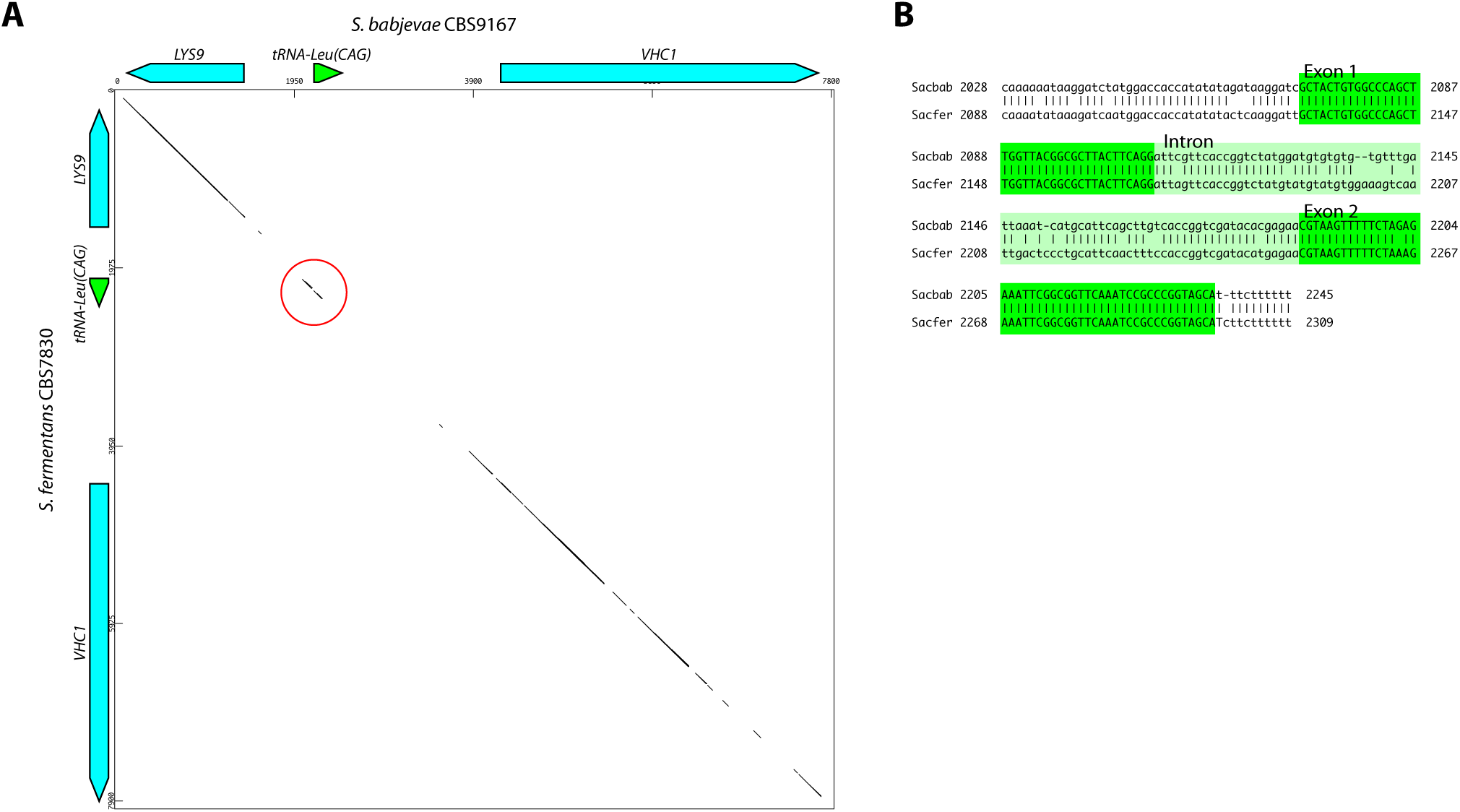
Conservation of the tRNA-Leu(CAG) locus between *S. babjevae* and *S. fermentans*. (A) Dot matrix plot of the region containing the tRNA-Leu(CAG) locus and the neighboring protein-coding genes *LYS9* and *VHC1*. Dots indicate regions with ≥ 25 matches in a 30 bp sliding window. The match at the tRNA locus is circled. (B) BLASTN alignment corresponding to the region circled in A. The exons of the tRNA gene are better conserved (only 1 nt different, i.e. 98.8% identity) than the intron (77.8% identity) and the aligned region upstream of the gene (83.7% identity). Regions further upstream and downstream cannot be aligned, as shown by the dot matrix plot.

## Discussion

Our work has generated extensive data about the tRNA-Leu(CAG) gene in the CUG-Ser2 clade species, but the functional role of this enigmatic gene is still not fully resolved. There is evidence both to support the hypothesis that the gene is functional, and to support the hypothesis that it is not functional, as summarized below.

The observations that suggest that the tRNA-Leu(CAG) gene is functional are:

● It is used (simultaneously with tRNA-Ser(CAG)) for translation in *Ascoidea asiatica* (Muhlhausen et al. 2018).
● tRNA-Leu(CAG) (as well as tRNA-Ser(CAG)) is transcribed and aminoacylated in *S. malanga* (Shulgina and Eddy 2021).
● A candidate tRNA-Leu(CAG) gene can be identified in most *Saccharomycopsis* genomes (Fig. 1B).
● The gene is more conserved in sequence than the flanking DNA and its intron (Fig. 7).
● There is a strong bioinformatic signal suggesting that some CUG codons are translated as leucine during sporulation (Fig. 3).

The observations that suggest that the tRNA-Leu(CAG) gene is not functional are:

● No translation of CUG codons as leucine has been detected in any of the five *Saccharomycopsis* species that have been examined by mass spectrometry (*S. capsularis*, *S. malanga*, *S. fibuligera*, *S. schoenii*, and *S. fermentans*) ((Krassowski et al. 2018; Muhlhausen et al. 2018; Junker et al. 2019) and this study).
● The gene is completely absent in a few species (*S. microspora*, *S. synnaedendra*, and *A. rubescens*) (Fig. 1B).
● In *Saccharomycopsis* species, there are numerous mutations in the gene’s sequence that alter bases that are usually very highly conserved in tRNA-Leu molecules, or that alter the stems and loops of the cloverleaf structure in unprecedented ways (Fig. 5; Fig. 6).

To reconcile these conflicting observations, we suggest that the tRNA-Leu(CAG) gene is dying in the CUG-Ser2 clade. Its role in translation of CUG codons is being replaced by tRNA-Ser(CAG), and this process of replacement has proceeded to different extents in different species within the CUG-Ser2 clade. It has proceeded further in *Saccharomycopsis* and in *A. rubescens* than in *A. asiatica.* The bioinformatic signal of evolutionary conservation of CUG-Leu translation in sporulation genes that we detected in the clade may reflect the fact that sporulation is a rare part of the life cycle and does not occur in every generation (Tsai et al. 2008), so the selection pressure to replace CUG codons in sporulation genes may be less strong than in other genes, causing them to be replaced more slowly. It is possible that such a bioinformatic signal could remain detectable in *Saccharomycopsis* genomes even millions of years after all translation of CUG codons as Leu has stopped.

We also suggest that the tRNA-Leu(CAG) gene may have an unknown non-translational function, that has led to the evolutionary retention of the gene in most species while allowing its sequence to diverge to a greater extent than is normal for a tRNA gene. If this non-translational function requires transcription of the locus, it can explain the transcription and splicing that has been observed (in *S. malanga* and *S. capsularis*, by RT-PCR or Northern blotting (Krassowski et al. 2018; Shulgina and Eddy 2021)), and it can explain the greater conservation of the exons than the intron (Fig. 7). If the non-translational function also requires leucylation of the transcript, it can explain the aminoacylation that was detected in *S. malanga* (Shulgina and Eddy 2021).

Many non-translational roles have been identified for tRNAs (Avcilar-Kucukgoze and Kashina 2020). An interesting precedent for what may have happened to the yeast tRNA-Leu(CAG) has been found in a glutamate tRNA gene in plant chloroplast genomes. As well as its role in translation of GAA codons, chloroplast tRNA-Glu(UUC) is also required for biosynthesis of mitochondrial heme (Howe and Smith 1991; Barbrook et al. 2006). The first step of heme synthesis uses as a substrate a glutamate residue, which must be attached (charged) to tRNA-Glu(UUC). In the non-photosynthetic parasitic plant *Balanophora*, which appears not to have a functioning native chloroplast translation apparatus and has lost all other tRNA genes, a degenerated tRNA-Glu(UUC) gene remains in its chloroplast genome (Su et al. 2019). The gene is badly damaged and lacks a normal cloverleaf, but the acceptor stem region is intact. It has been proposed that this stem is aminoacylated with glutamate, which is then used for heme synthesis, and that the gene has no translational function (Su et al. 2019). Notably, the acceptor stem of tRNA-Leu(CAG) is well conserved in *Saccharomycopsis* (Fig. 5; Fig. 6).

Additionally, in both eukaryotes and prokaryotes, charged tRNAs are used as a source of amino acids by a family of enzymes termed aminoacyl-tRNA protein transferases, such as Ate1 which catalyses the arginylation of proteins in a tRNA-dependent but ribosome-independent manner (Abeywansha et al. 2023). Ate1 is conserved from yeast to mammals and is essential for embryogenesis in mice (Saha and Kashina 2011) and has been implicated in the Arg/N-end rule degron pathway (Oh et al. 2020). A similar mechanism exists in Gram negative bacteria, where the enzyme Leu/Phe transferase tags proteins for degradation by leucylation or phenylation (Tobias et al. 1991; Shrader et al. 1993). These two groups of aminoacyl-tRNA protein transferases are believed to have evolved independently (Suto et al. 2006). We speculate that *Saccharomycopsis* tRNA-Leu(CAG) may have an analogous non-translational function, similar to one of these pathways, that requires its transcription, splicing and aminoacylation.

## Materials and Methods

### Genome sequencing

We sequenced 23 strains of *Saccharomycopsis* by Illumina technology, including the type strains of most of the known species in the genus (Table S1). Strains were purchased from the Westerdijk Institute for Fungal Diversity (CBS Collection, The Netherlands). Cultures of all strains were grown in YPD at 30°C overnight. Cells were harvested by centrifugation and cell pellets were resuspended in 200 µl extraction buffer (2% Triton X-100, 100 mM NaCl, 10 mM Tris pH 7.4, 1 mM EDTA, 1% SDS) in a 1.5 ml screw-cap tube. Approximately 0.3 g acid-washed glass beads (425-600 μm) were added with 200 μl phenol/chloroform/isoamyl alcohol (25:24:1). The mixture was agitated on a 600 MiniG bead beater (Spex SamplePrep) at 1,500 rpm, 4-6 times for 30 s each, and centrifuged at 15,000 rcf for 10 min. The top aqueous layer was transferred to a new 1.5 ml screw-cap tube, 200 μl TE buffer was added, and 200 μl of the phenol/chloroform/isoamyl alcohol mixture was added. This was agitated as before on the 600 MiniG bead beater, centrifuged at 14,000 rpm for 10 min. The top aqueous layer was transferred to a new microfuge tube, after which 80 μl 7.5 M ammonium acetate and 1 ml 100% isopropyl alcohol were added to precipitate the DNA. DNA was pelleted by centrifugation at 14,000 rpm, washed using 70% ethanol and dried in a SpeedVac (Eppendorf Concentrator 5301 at 45°C for 2 min pulses until dry). Pellets were resuspended in 400 µl TE buffer with 1 µl RNase A (10 mg/ml) and incubated overnight at 37°C. DNA was re-precipitated and washed once more as above and re-suspended in 150 µl water. DNA quality and concentration was assessed by gel electrophoresis, Nanodrop, and Qubit measurement. Genomic DNA was sequenced by BGI Tech Solutions (Hong Kong) using an Illumina X Ten instrument (150 bp paired ends, 1.5 Gb raw data per sample).

### Genome assembly, annotation and phylogenomics

Paired end reads for the 23 strains of this study were quality assessed by FASTQC before and after trimming by Skewer v.0.2.2 (Jiang et al. 2014). *De novo* genome assemblies were generated by SPAdes v.3.11 (Bankevich et al. 2012) for each set of reads. Genome quality was assessed by QUAST and coverage-versus-length plots (Douglass et al. 2019). Ten other genome sequences were downloaded from NCBI (Table S1). For each of the 33 genome sequences, AUGUSTUS v3.5.0 (Stanke et al. 2008) was used to predict ORFs, and tRNAscan-SE v2.0.5 (Lowe and Eddy 1997) was used to annotate tRNA genes. For some *Saccharomycopsis* species, the tRNA-Leu(CAG) gene was not identified by tRNAscan-SE and instead was annotated manually based on BLASTN search results. All predicted tRNA genes with CAG as the anticodon were manually inspected and classified as either Ser or Leu tRNAs using identity determinants specific to tRNA-Leu or tRNA-Ser molecules (Giegé et al. 1998) as well as synteny conservation.

For phylogenomic analysis, the predicted sets of ORFs from Augustus were trimmed to remove ORFs that were incomplete because they reached the end of a contig. This set of ORFs was translated using the standard genetic code, except that CUG codons were translated as ‘X’. These 33 sets of translated ORFs from each genome were used as input to OrthoFinder v2.5.2 (Emms and Kelly 2019). The resulting set of all single copy orthologs (1,226) was used for phylogenomic tree construction. These orthogroups were each aligned using MUSCLE v3.8.31 (Edgar 2004), trimmed using trimAl v1.2 (Capella-Gutierrez et al. 2009) with the -automatic heuristic setting, and concatenated together into a supermatrix containing 567,098 sites. IQ-tree v1.6.12 (Nguyen et al. 2015) was used to first find an appropriate maximum likelihood model for this supermatrix, and then to construct a tree using that model (LG+F+R5), performing 1000 ultrafast bootstraps and 1000 sh-alrt tests. The branching order of the tree was inspected manually and compared to the OrthoFinder species tree, which is inferred from gene duplication events.

### Computational search for conserved CUG codons and orthogroup homolog assignment

Orthologous protein groups (orthogroups) that contained at least 25 single copy orthologs (4,187 total groups) were used for CUG conservation analysis. Multi-copy orthologs were discarded so that only single copy genes were used in the downstream analysis. Each orthogroup was aligned using MUSCLE and trimmed using trimAl as described above, to remove poorly conserved columns. Trimmed alignments were then analyzed as described in the text and Figure 3 to generate Column Scores for each column and an overall CUG-Leu Conservation Score for the orthogroup. To identify *S. cerevisiae* orthologs of each orthogroup, bidirectional BLASTP searches were carried out between the *S. cerevisiae* proteome (Engel et al. 2022) and the predicted proteome of each sequenced genome, to identical reciprocal best hits (RBHs) for each genome. To assign an *S. cerevisiae* gene name to an orthogroup, we required it to be the RBH of the majority of the 25-33 genes in the orthogroup.

### CRISPR RNP gene editing in *S. capsularis*

*S. capsularis* Y-17639 was chosen for gene editing attempts by electroporation. We adapted a CRISPR RNP protocol from Grahl et al. (2017) and DiCarlo et al. (2013). *S. capsularis* NRRL Y-17639 was inoculated in 5 ml YPD, grown at 30°C overnight, back-diluted to OD_600_ 0.1 in 50 ml YPD, and grown for 14 h at 30°C. Cells were harvested by centrifugation at 1780 rcf at 4°C. The cells were washed in 10 ml ice-cold sterile double-deionised water, centrifuged as before, and washed again in 25 ml ice-cold electroporation buffer (EB; 1 M sorbitol, 1 mM CaCl_2_). Cells were then centrifuged, resuspended in lithium acetate/DTT (500 mM/10 mM), and incubated for 30 m at 30°C. They were then centrifuged, washed a final time in 25 ml EB, and resuspended in 200 μl EB. They were kept on ice until needed, and 40 μl of this mixture was used for each transformation reaction.

crRNA and tracrRNA ssRNA oligos from the ALT-R system (Integrated DNA Technologies) were annealed as recommended. The annealed guides were incubated with Cas9p for 5 min at room temperature, and then kept on ice until ready. For homology directed repair templates, ‘CUGless’ drug marker cassettes with no CUG codons were amplified from plasmid stocks with 70-mer oligonucleotide primers, with 20 bases matching the ends of the marker and 50 bases matching either directly adjacent to the CRISPR cut site (Fig. S1), or several hundred bases upstream/downstream of the cut site (Fig. 2). Typically 4 × 50 μl PCR reactions were pooled together, purified by spin columns, and eluted in 30 μl water to generate each repair template. The CRISPR RNPs (6.6 μl), 2.5 μg of repair template DNA, and 40 μl of the prepared competent cells were mixed together in a microfuge tube on ice. This mixture was transferred into a 2 mm sterile electroporation cuvette (VWR) chilled on ice, and electroporated (Bio-Rad GenePulser Xcell; 2,500 V, 25µF, 200Ω). The electroporated PCR mixture was transferred to 7 ml recovery medium (1:1 YPD:1 M sorbitol), and incubated at 30°C for 2.5 h. Cells were then spun down, resuspended in 200 μl YPD, and plated to the relevant drugs. For selection, 50 ng/μl G-418 or 10 ng/μl nourseothricin was used.

Correct integration of drug cassettes at the target locus was assessed first by colony PCR of the integration junctions. For colony PCR in *S. capsularis*, we developed a rapid protocol using phenol/chloroform/isoamyl alcohol. Briefly, a colony is resuspended in 200 μl TE, 200 μl phenol/chloroform/isoamyl alcohol was added with approximately 0.3 g acid-washed glass beads (425-600 μm), all in a 1.5 ml screw-cap tube. The whole mixture is treated in a BeadBeater 4-6 times for 30 s each, then centrifuged at 15,000 rcf. The top aqueous layer is transferred to a new tube. This TE layer, although phenol contaminated, can be used for many PCR reactions (0.5-1 μl per 20-50 μl PCR reaction) and frozen for future use. Once each integration junction was detected by PCR, the entire locus was amplified by PCR using external primers and sent for Sanger sequencing to confirm its structure. Sequences of oligonucleotides used as PCR primers are given in Table S2.

### Tandem Mass Spectrometry

All strains were grown on either sporulation media (YM agar, 25°C, 24 days for all *S. capsularis* strains; 5% malt extract agar, 25°C, 25 days for *S. fermentans*) or vegetative media (YPD agar, 25°C, 3 days). Cells were scraped from the plates, resuspended in sterile water, and brightfield images were taken to verify sporulation.

For *S. capsularis*, samples were processed using all reagents and instructions supplied in the Preomics iST and iST Fractionation add-on kits. In brief, replicate cell pellets for each sample underwent one of two treatments: one set was subjected to a column-based fractionation (Preomics iST Fractionation kit) and the other set were treated as whole cell lysates. Cell pellets were solubilized in 150 μL of “Lyse” buffer (containing Tris-HCl, sodium deoxycholate (SDC), 0.1% sodium dodecyl sulfate (SDS), tris(2-carboxyethyl)phosphine (TCEP), and 2-chloroacetamide) and heated to 95 °C for 10 min. Next, 50 μL of the resulting denatured, reduced, and alkylated solution was transferred to the reaction vessel. Enzyme (LysC and trypsin) was added, and samples were hydrolyzed at 37 °C for 3 h. The resulting peptide mixture was washed and eluted as per the manufacturer’s instructions. The digested peptides were fractionated as per manufacturer’s instructions before subsequently vacuum-dried and dissolved in water/0.5% acetic acid.

The samples were analyzed by the Mass Spectrometry Resource (MSR) in University College Dublin on a Thermo Scientific Q Exactive mass spectrometer connected to a Dionex Ultimate 3000 (RSLCnano) chromatography system. Peptides were separated on C18 home-made column (C18-AQ Dr. Maisch Reprosil-Pur 100 × 0.075 mm × 3 μm) over 60 min (for pseudo-fractions) or 180 min (for whole cell lysates) at a flow rate of 250 nL/min with a linear gradient of increasing ACN from 1% to 27%. The mass spectrometer was operated in data dependent mode; a high resolution (70,000) MS scan (300-1600 m/z) was performed to select the twelve most intense ions and fragmented using high energy C-trap dissociation for MS/MS analysis.

For *S. fermentans*, total protein extract was run on an SDS-PAGE gel and cut into approximately 10 fractions for both vegetative and sporulative cells. In-gel digestion with trypsin was performed on each excised fraction, using the protocol of Shevchenko et al. {, 2006 #7505} with minor changes. Briefly, de-stained gel pieces were reduced with 10 mM DTT for 45 min at 56°C and alkylated with 55 mM iodoacetamide in the dark for 30 min at room temperature. All reactions were performed in 50 mM ammonium bicarbonate buffer and gel pieces were dehydrated between steps. Dehydrated gel pieces were rehydrated in trypsin solution (20 ng/μl trypsin, 50 mM ammonium bicarbonate) on ice for 45 min followed by overnight incubation at 37°C. Tryptic peptides were extracted with two rounds of extraction buffer (100 μl; 70% acetonitrile, 5% formic acid) pooled extracts were dried under vacuum and resuspended in 0.1% formic acid. Εxtracted peptides were loaded onto C18 EvoTip disposable trap columns and run on a timsTOF Pro mass spectrometer (Bruker Daltonics, Bremen, Germany) coupled to the EvoSep One system (EvoSep, Odense, Denmark). The peptides were separated on a reversed-phase C18 Endurance column (15 cm × 150 μm ID, C 18, 1.9 μm, EV-1106) using the pre-set Extended 15 SPD method. Mobile phases were 0.1% (v/v) formic acid in water (phase A) and 0.1% (v/v) formic acid in acetonitrile (phase B). The peptides were separated by an increasing gradient of mobile phase B at 220 nL/min for 88 minutes. The timsTOF Pro mass spectrometer was operated in positive ion polarity with TIMS (Trapped Ion Mobility Spectrometry) and PASEF (Parallel Accumulation Serial Fragmentation) modes enabled. The accumulation and ramp times for the TIMS were both set to 100 ms, with an ion mobility (1/k0) range from 0.6 to 1.60 V.s/cm^2^. Spectra were recorded in the mass range from 100 to 1,700 m/z. The precursor (MS) Intensity Threshold was set to 2,500 and the precursor Target Intensity set to 20,000. Each PASEF cycle consisted of one MS ramp for precursor detection followed by 10 PASEF MS/MS ramps, with a total cycle time of 1.16 s.

### Proteomic data processing and bioinformatics

Raw data from the Q-Exactive and timsTOF was processed using MaxQuant (Cox and Mann 2008; Tyanova et al. 2016) (version 2.0.3.0) incorporating the Andromeda search engine (Cox et al. 2011). To identify peptides, the MS/MS spectra for each strain for each condition were jointly run in a database search; the spectra were matched against 19 custom FASTA databases specific to each strain’s genome; each possible translation of CUG excluding isoleucine accounts for the 19 databases. All searches were performed using the default setting of MaxQuant, with trypsin as specified enzyme allowing two missed cleavages and a false discovery rate of 1% on the peptide and protein level. The database searches were performed with carbamidomethyl (C) as fixed modification and acetylation (protein N terminus) and oxidation (M) as variable modifications. Codon counts of all amino acid translations of CUG were plotted using R.

## Data availability

New genome sequences reported in this study are listed in Table S1 and have been deposited in the ENA/NCBI/DDBJ databases under BioProject PRJNA977123.

## Acknowledgements

This work was supported by the European Research Council (789341). We thank Padraic Heneghan for discussion.

## References

Abeywansha T, Huang W, Ye X, Nawrocki A, Lan X, Jankowsky E, Taylor DJ, Zhang Y. 2023. The structural basis of tRNA recognition by arginyl-tRNA-protein transferase. Nat Commun 14:2232.

Avcilar-Kucukgoze I, Kashina A. 2020. Hijacking tRNAs From Translation: Regulatory Functions of tRNAs in Mammalian Cell Physiology. Front Mol Biosci 7:610617.

Bankevich A, Nurk S, Antipov D, Gurevich AA, Dvorkin M, Kulikov AS, Lesin VM, Nikolenko SI, Pham S, Prjibelski AD, et al. 2012. SPAdes: a new genome assembly algorithm and its applications to single-cell sequencing. J Comput Biol 19:455–477.

Barbrook AC, Howe CJ, Purton S. 2006. Why are plastid genomes retained in non-photosynthetic organisms? Trends Plant Sci 11:101–108.

Boyle EI, Weng S, Gollub J, Jin H, Botstein D, Cherry JM, Sherlock G. 2004. GO::TermFinder--open source software for accessing Gene Ontology information and finding significantly enriched Gene Ontology terms associated with a list of genes. Bioinformatics 20:3710–3715.

Brar GA, Yassour M, Friedman N, Regev A, Ingolia NT, Weissman JS. 2012. High-resolution view of the yeast meiotic program revealed by ribosome profiling. Science 335:552–557.

Capella-Gutierrez S, Silla-Martinez JM, Gabaldon T. 2009. trimAl: a tool for automated alignment trimming in large-scale phylogenetic analyses. Bioinformatics 25:1972–1973.

Choo JH, Hong CP, Lim JY, Seo JA, Kim YS, Lee DW, Park SG, Lee GW, Carroll E, Lee YW, et al. 2016. Whole-genome de novo sequencing, combined with RNA-Seq analysis, reveals unique genome and physiological features of the amylolytic yeast *Saccharomycopsis fibuligera* and its interspecies hybrid. Biotechnol Biofuels 9:246.

Chu S, DeRisi J, Eisen M, Mulholland J, Botstein D, Brown PO, Herskowitz I. 1998. The transcriptional program of sporulation in budding yeast. Science 282:699–705.

Cox J, Mann M. 2008. MaxQuant enables high peptide identification rates, individualized p.p.b.-range mass accuracies and proteome-wide protein quantification. Nat Biotechnol 26:1367–1372.

Cox J, Neuhauser N, Michalski A, Scheltema RA, Olsen JV, Mann M. 2011. Andromeda: a peptide search engine integrated into the MaxQuant environment. J Proteome Res 10:1794–1805.

Crick FH. 1968. The origin of the genetic code. J Mol Biol 38:367–379.

Dicarlo JE, Norville JE, Mali P, Rios X, Aach J, Church GM. 2013. Genome engineering in *Saccharomyces cerevisiae* using CRISPR-Cas systems. Nucleic Acids Res 41:4336–4343.

Douglass AP, O’Brien CE, Offei B, Coughlan AY, Ortiz-Merino RA, Butler G, Byrne KP, Wolfe KH. 2019. Coverage-Versus-Length Plots, a Simple Quality Control Step for de Novo Yeast Genome Sequence Assemblies. G3 (Bethesda) 9:879–887.

Edgar RC. 2004. MUSCLE: a multiple sequence alignment method with reduced time and space complexity. BMC Bioinformatics 5:113.

Emms DM, Kelly S. 2019. OrthoFinder: phylogenetic orthology inference for comparative genomics. Genome Biol 20:238.

Engel SR, Wong ED, Nash RS, Aleksander S, Alexander M, Douglass E, Karra K, Miyasato SR, Simison M, Skrzypek MS, et al. 2022. New data and collaborations at the *Saccharomyces* Genome Database: updated reference genome, alleles, and the Alliance of Genome Resources. Genetics 220.

Giegé R, Sissler M, Florentz C. 1998. Universal rules and idiosyncratic features in tRNA identity. Nucleic Acids Res 26:5017–5035.

Grahl N, Demers EG, Crocker AW, Hogan DA. 2017. Use of RNA-Protein Complexes for Genome Editing in Non-albicans Candida Species. mSphere 2.

Howe CJ, Smith AL. 1991. Plants without chlorophyll. Nature 349:109.

Jacques N, Louis-Mondesir C, Coton M, Coton E, Casaregola S. 2014. Two novel Saccharomycopsis species isolated from black olive brines and a tropical plant. Description of Saccharomycopsis olivae f. a., sp. nov. and Saccharomycopsis guyanensis f. a., sp. nov. Reassignment of Candida amapae to Saccharomycopsis amapae f. a., comb. nov., Candida lassenensis to Saccharomycopsis lassenensis f. a., comb. nov. and Arthroascus babjevae to Saccharomycopsis babjevae f. a., comb. nov. Int J Syst Evol Microbiol 64:2169–2175.

Jiang H, Lei R, Ding SW, Zhu S. 2014. Skewer: a fast and accurate adapter trimmer for next-generation sequencing paired-end reads. BMC Bioinformatics 15:182.

Junker K, Chailyan A, Hesselbart A, Forster J, Wendland J. 2019. Multi-omics characterization of the necrotrophic mycoparasite Saccharomycopsis schoenii. PLoS Pathog 15:e1007692.

Kawaguchi Y, Honda H, Taniguchi-Morimura J, Iwasaki S. 1989. The codon CUG is read as serine in an asporogenic yeast *Candida cylindracea*. Nature 341:164–166.

Keeling PJ. 2016. Genomics: Evolution of the genetic code. Curr Biol 26:R851–853.

Kollmar M, Mühlhausen S. 2017a. How tRNAs dictate nuclear codon reassignments: Only a few can capture non-cognate codons. RNA Biol 14:293–299.

Kollmar M, Mühlhausen S. 2017b. Nuclear codon reassignments in the genomics era and mechanisms behind their evolution. Bioessays 39:1600221.

Krassowski T, Coughlan AY, Shen XX, Zhou X, Kominek J, Opulente DA, Riley R, Grigoriev IV, Maheshwari N, Shields DC, et al. 2018. Evolutionary instability of CUG-Leu in the genetic code of budding yeasts. Nat Commun 9:1887.

Kreger-van Rij NJW. 1964. A taxonomic study of the yeast genera Endomycopsis, Pichia and Debaryomyces. [PhD]: PhD thesis, Leiden University.

Kuleshov MV, Diaz JEL, Flamholz ZN, Keenan AB, Lachmann A, Wojciechowicz ML, Cagan RL, Ma’ayan A. 2019. modEnrichr: a suite of gene set enrichment analysis tools for model organisms. Nucleic Acids Res 47:W183–W190.

Kurtzman CP, Fell JW, Boekhout T. 2011. The Yeasts, a Taxonomic Study. In. Amsterdam: Elsevier.

Kurtzman CP, Smith MT. 2011. *Saccharomycopsis* Schiönning (1903). In: Kurtzman CP, Fell JW, Boekhout T, editors. The Yeasts, A Taxonomic Study. Amsterdam: Elsevier. p. 751–763.

Lin BY, Chan PP, Lowe TM. 2019. tRNAviz: explore and visualize tRNA sequence features. Nucleic Acids Res 47:W542–W547.

Lodder J, Kreger-van Rij NJW. 1952. The Yeasts, A Taxonomic Study. Amsterdam: North-Holland Publishing Company.

Lowe TM, Eddy SR. 1997. tRNAscan-SE: a program for improved detection of transfer RNA genes in genomic sequence. Nucleic Acids Res 25:955–964.

Minh BQ, Schmidt HA, Chernomor O, Schrempf D, Woodhams MD, von Haeseler A, Lanfear R. 2020. IQ-TREE 2: New Models and Efficient Methods for Phylogenetic Inference in the Genomic Era. Mol Biol Evol 37:1530–1534.

Mordret E, Dahan O, Asraf O, Rak R, Yehonadav A, Barnabas GD, Cox J, Geiger T, Lindner AB, Pilpel Y. 2019. Systematic Detection of Amino Acid Substitutions in Proteomes Reveals Mechanistic Basis of Ribosome Errors and Selection for Translation Fidelity. Mol Cell 75:427–441 e425.

Mühlhausen S, Findeisen P, Plessmann U, Urlaub H, Kollmar M. 2016. A novel nuclear genetic code alteration in yeasts and the evolution of codon reassignment in eukaryotes. Genome Res 26:945–955.

Muhlhausen S, Schmitt HD, Pan KT, Plessmann U, Urlaub H, Hurst LD, Kollmar M. 2018. Endogenous Stochastic Decoding of the CUG Codon by Competing Ser- and Leu-tRNAs in *Ascoidea asiatica*. Curr Biol 28:2046–2057 e2045.

Neiman AM. 2011. Sporulation in the budding yeast *Saccharomyces cerevisiae*. Genetics 189:737–765.

Nguyen LT, Schmidt HA, von Haeseler A, Minh BQ. 2015. IQ-TREE: a fast and effective stochastic algorithm for estimating maximum-likelihood phylogenies. Mol Biol Evol 32:268–274.

Oh JH, Hyun JY, Chen SJ, Varshavsky A. 2020. Five enzymes of the Arg/N-degron pathway form a targeting complex: The concept of superchanneling. Proc Natl Acad Sci U S A 117:10778–10788.

Osawa S. 1995. Evolution of the Genetic Code. Oxford: Oxford University Press.

Prat L, Heinemann IU, Aerni HR, Rinehart J, O’Donoghue P, Soll D. 2012. Carbon source-dependent expansion of the genetic code in bacteria. Proc Natl Acad Sci U S A 109:21070–21075.

Quintilla R, Kolecka A, Casaregola S, Daniel HM, Houbraken J, Kostrzewa M, Boekhout T, Groenewald M. 2018. MALDI-TOF MS as a tool to identify foodborne yeasts and yeast-like fungi. Int J Food Microbiol 266:109–118.

Riley R, Haridas S, Wolfe KH, Lopes MR, Hittinger CT, Goker M, Salamov AA, Wisecaver JH, Long TM, Calvey CH, et al. 2016. Comparative genomics of biotechnologically important yeasts. Proc Natl Acad Sci U S A 113:9882–9887.

Rother M, Krzycki JA. 2010. Selenocysteine, pyrrolysine, and the unique energy metabolism of methanogenic archaea. Archaea 2010.

Saha S, Kashina A. 2011. Posttranslational arginylation as a global biological regulator. Dev Biol 358:1–8.

Santos MA, Tuite MF. 1995. The CUG codon is decoded *in vivo* as serine and not leucine in *Candida albicans*. Nucleic Acids Res 23:1481–1486.

Sekulovski S, Trowitzsch S. 2022. Transfer RNA processing - from a structural and disease perspective. Biol Chem 403:749–763.

Shimodaira H. 2002. An approximately unbiased test of phylogenetic tree selection. Syst Biol 51:492–508.

Shrader TE, Tobias JW, Varshavsky A. 1993. The N-end rule in Escherichia coli: cloning and analysis of the leucyl, phenylalanyl-tRNA-protein transferase gene aat. J Bacteriol 175:4364–4374.

Shulgina Y, Eddy SR. 2021. A computational screen for alternative genetic codes in over 250,000 genomes. Elife 10.

Stanke M, Diekhans M, Baertsch R, Haussler D. 2008. Using native and syntenically mapped cDNA alignments to improve de novo gene finding. Bioinformatics 24:637–644.

Su HJ, Barkman TJ, Hao W, Jones SS, Naumann J, Skippington E, Wafula EK, Hu JM, Palmer JD, dePamphilis CW. 2019. Novel genetic code and record-setting AT-richness in the highly reduced plastid genome of the holoparasitic plant Balanophora. Proc Natl Acad Sci U S A 116:934–943.

Subramanian A, Tamayo P, Mootha VK, Mukherjee S, Ebert BL, Gillette MA, Paulovich A, Pomeroy SL, Golub TR, Lander ES, et al. 2005. Gene set enrichment analysis: a knowledge-based approach for interpreting genome-wide expression profiles. Proc Natl Acad Sci U S A 102:15545–15550.

Sugita T, Nakase T. 1999. Non-universal usage of the leucine CUG codon and the molecular phylogeny of the genus *Candida*. Syst Appl Microbiol 22:79–86.

Suto K, Shimizu Y, Watanabe K, Ueda T, Fukai S, Nureki O, Tomita K. 2006. Crystal structures of leucyl/phenylalanyl-tRNA-protein transferase and its complex with an aminoacyl-tRNA analog. EMBO J 25:5942–5950.

Tobias JW, Shrader TE, Rocap G, Varshavsky A. 1991. The N-end rule in bacteria. Science 254:1374–1377.

Tsai IJ, Bensasson D, Burt A, Koufopanou V. 2008. Population genomics of the wild yeast *Saccharomyces paradoxus*: quantifying the life cycle. Proc Natl Acad Sci U S A 105:4957–4962.

Tyanova S, Temu T, Cox J. 2016. The MaxQuant computational platform for mass spectrometry-based shotgun proteomics. Nat Protoc 11:2301–2319.

Xie Z, Bailey A, Kuleshov MV, Clarke DJB, Evangelista JE, Jenkins SL, Lachmann A, Wojciechowicz ML, Kropiwnicki E, Jagodnik KM, et al. 2021. Gene Set Knowledge Discovery with Enrichr. Curr Protoc 1:e90.

Yuan J, O’Donoghue P, Ambrogelly A, Gundllapalli S, Sherrer RL, Palioura S, Simonovic M, Soll D. 2010. Distinct genetic code expansion strategies for selenocysteine and pyrrolysine are reflected in different aminoacyl-tRNA formation systems. FEBS Lett 584:342–349.

Yuan X, Peng K, Li C, Zhao Z, Zeng X, Tian F, Li Y. 2021. Complete Genomic Characterization and Identification of *Saccharomycopsis phalluae* sp. nov., a Novel Pathogen Causes Yellow Rot Disease on *Phallus rubrovolvatus*. J Fungi (Basel) 7.

