## Supplementary Information for "A genetic code change in progress: tRNA-Leu(CAG) is conserved in most *Saccharomycopsis* yeast species but is non-essential and does not compete with tRNA-Ser(CAG) in translation"

### Supplementary Figures

**Figure S1**. Disruption of the *ADE2* gene of *S. capsularis* NRRL Y-17639 using CRISPR-Cas9 ribonucleoproteins.
(A) Schematic of construct design. DNA elements are drawn to scale, and numbers denote distance in basepairs. The dashed red line denotes the CRISPR-Cas9 RNP cut site, and the red box above it represents the complex itself, assembled *in vitro* prior to transformation. The *pTEF1::KanR::tCYC1* construct used to disrupt *ADE2* was amplified by PCR using primers EOC17 and EOC18 from a template containing a CUGless version of the KanMX cassette. Locations of primers used in the screening of transformants are shown. The *ADE2* gene sequence of *S. capsularis* NRRL Y-17639 was taken from NCBI accession number PPIG01000008.1.
(B) A representative PCR screen of 30 independent transformants from two replicate transformations using the KanMX construct as in (A). Primer pairs EOC1/EOC7 and EOC3/EOC8 were used to verify the upstream and downstream integration junctions, respectively.
(C) Streaks of wildtype (left) and a successful *ade2* knockout (right) to SC (synthetic complete) agar. A pinkish color is visible in the transformed strain.
(D) Colony screens of KanMX transformation (as depicted in A and B) and parallel transformations using CUGless nourseothricin (NTC) or hygromycin (Hyg) drug resistance cassettes to disrupt *ADE2,* patched to both YPD agar and SC-ADE agar (synthetic complete media lacking adenine). No growth is visible on the SC-ADE plates.

**Figure S2.** YeastEnrichr and Gene Set Enrichment Analysis of Gene Ontology analysis of genes with the top 5% of CUG-Leu Conservation Scores.

(A) Selected YeastEnrichr outputs of enriched GO terms in the top 5%.

(B) Volcano plot of Gene Set Enrichment Analysis (GSEA) of the top 5%. Each point is a GO term, plotted to show its enrichment score (X-axis) and statistical significance (Y-axis; False Discovery Rate). The color gradient indicates the number of genes contributing to each term.

**Figure S3.** Conservation of the tRNA-Leu(CAG) and tRNA-Ser(CAG) genes in the CUG-Ser2 clade. (A) Consensus and occupancy summary graphs of alignments of tRNA-Leu(CAG) genes (upper, 29 sequences) and tRNA-Ser(CAG) genes (lower, 41 sequences), from the CUG-Ser2 clade species analyzed in this study. At each position, the consensus plots indicate the proportion of sequences in the alignment that match the consensus (black bars indicate 100% conservation), and the occupancy plots indicate the proportion without an alignment gap. Sequences were aligned manually, and plots were made using the Jalview program (Waterhouse et al. 2009). (B) Unrooted phylogenetic trees built from the two alignments described above, using PhyML with default settings. The trees are to the same scale.

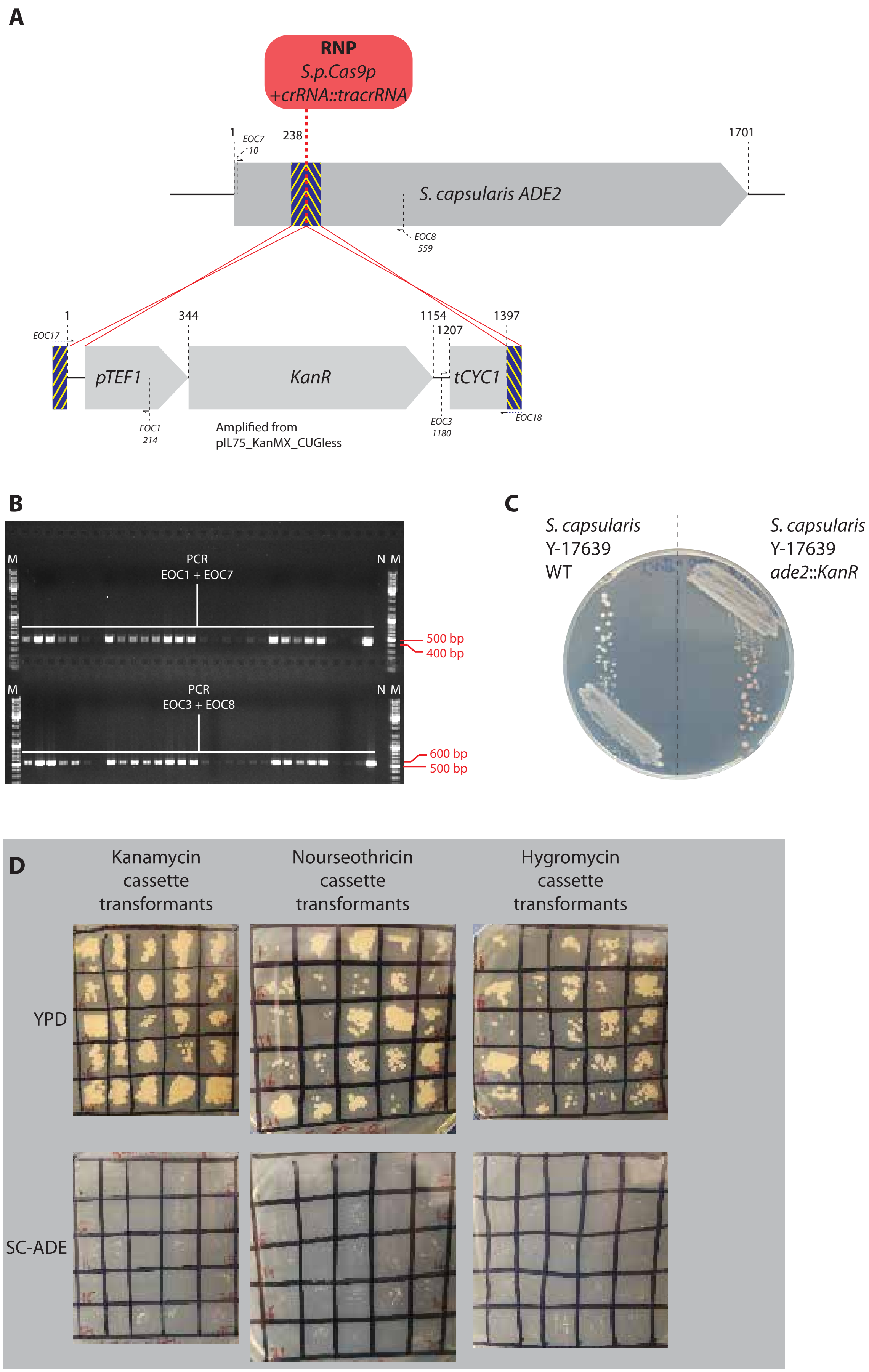
A
RNP
S.p.Cas9p
+crRNA::tracrRNA
1
EOC7
10
238
1701
S. capsularis ADE2
EOC8
559
1
344
1154
1207
1397
EOC17
pTEF1
KanR
tCYC1
EOC18
EOC1
214
Amplified from
pIL75_KanMX_CUGless
EOC3
1180
B
M
PCR
EOC1 + EOC7
N M
500 bp
400 bp
M
PCR
EOC3 + EOC8
N M
600 bp
500 bp
C
S. capsularis
Y-17639
WT
S. capsularis
Y-17639
ade2::KanR
D
Kanamycin
cassette
transformants
Nourseothricin
cassette
transformants
Hygromycin
cassette
transformants
YPD
SC-ADE

Figure S1

A

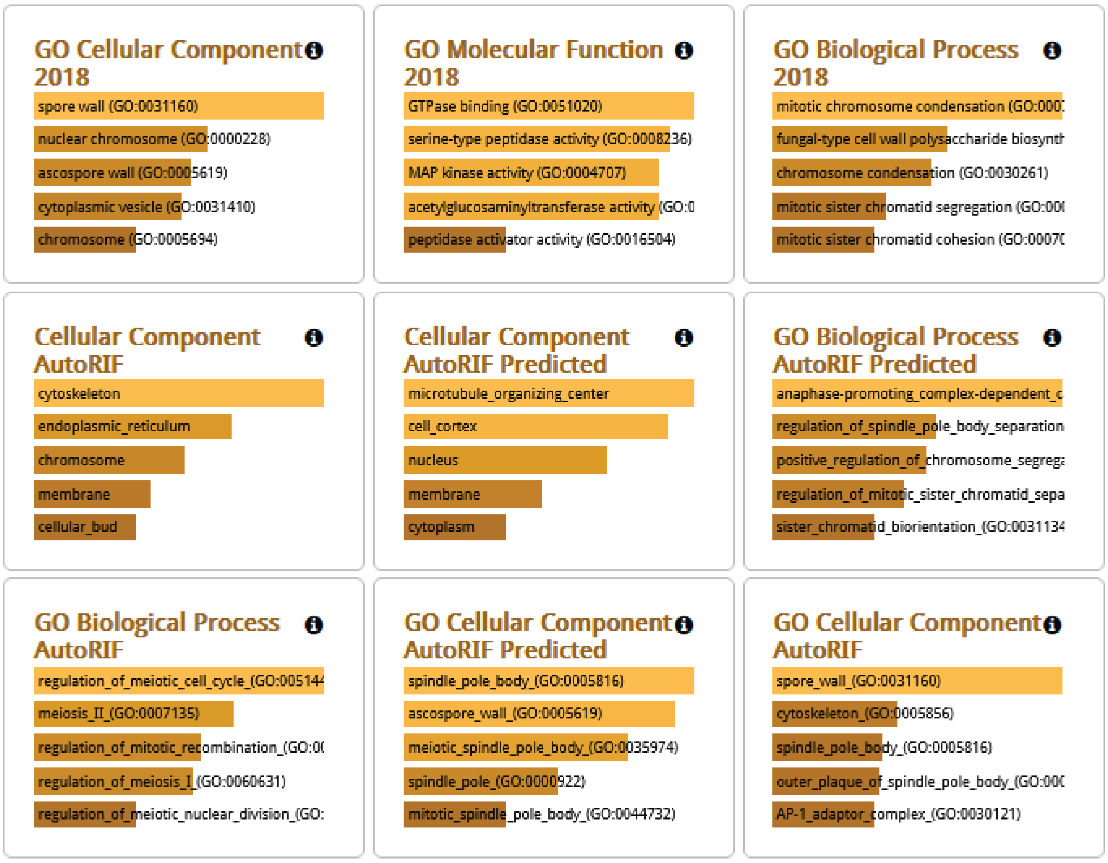
GO Cellular Component 2018
spore wall (GO:0031160)
nuclear chromosome (GO:0000228)
ascospore wall (GO:0005619)
cytoplasmic vesicle (GO:0031410)
chromosome (GO:0005694)
GO Molecular Function 2018
GTPase binding (GO:0051020)
serine-type peptidase activity (GO:0008236)
MAP kinase activity (GO:0004707)
acetylglucosaminyltransferase activity (GO:0
peptidase activator activity (GO:0016504)
GO Biological Process 2018
mitotic chromosome condensation (GO:000
fungal-type cell wall polysaccharide biosynth
chromosome condensation (GO:0030261)
mitotic sister chromatid segregation (GO:00
mitotic sister chromatid cohesion (GO:0007
Cellular Component AutoRIF
cytoskeleton
endoplasmic_reticulum
chromosome
membrane
cellular_bud
Cellular Component AutoRIF Predicted
microtubule_organizing_center
cell_cortex
nucleus
membrane
cytoplasm
GO Biological Process AutoRIF Predicted
anaphase-promoting_complex-dependent_c
regulation_of_spindle_pole_body_separation
positive_regulation_of_chromosome_segrega
regulation_of_mitotic_sister_chromatid_sepa
sister_chromatid_biorientation_(GO:0031134
GO Biological Process AutoRIF
regulation_of_meiotic_cell_cycle_(GO:005144
meiosis_II_(GO:0007135)
regulation_of_mitotic_recombination_(GO:0
regulation_of_meiosis_I_(GO:0060631)
regulation_of_meiotic_nuclear_division_(GO:
GO Cellular Component AutoRIF Predicted
spindle_pole_body_(GO:0005816)
ascospore_wall_(GO:0005619)
meiotic_spindle_pole_body_(GO:0035974)
spindle_pole_(GO:0000922)
mitotic_spindle_pole_body_(GO:0044732)
GO Cellular Component AutoRIF
spore_wall_(GO:0031160)
cytoskeleton_(GO:0005856)
spindle_pole_body_(GO:0005816)
outer_plaque_of_spindle_pole_body_(GO:000
AP-1_adaptor_complex_(GO:0030121)

B

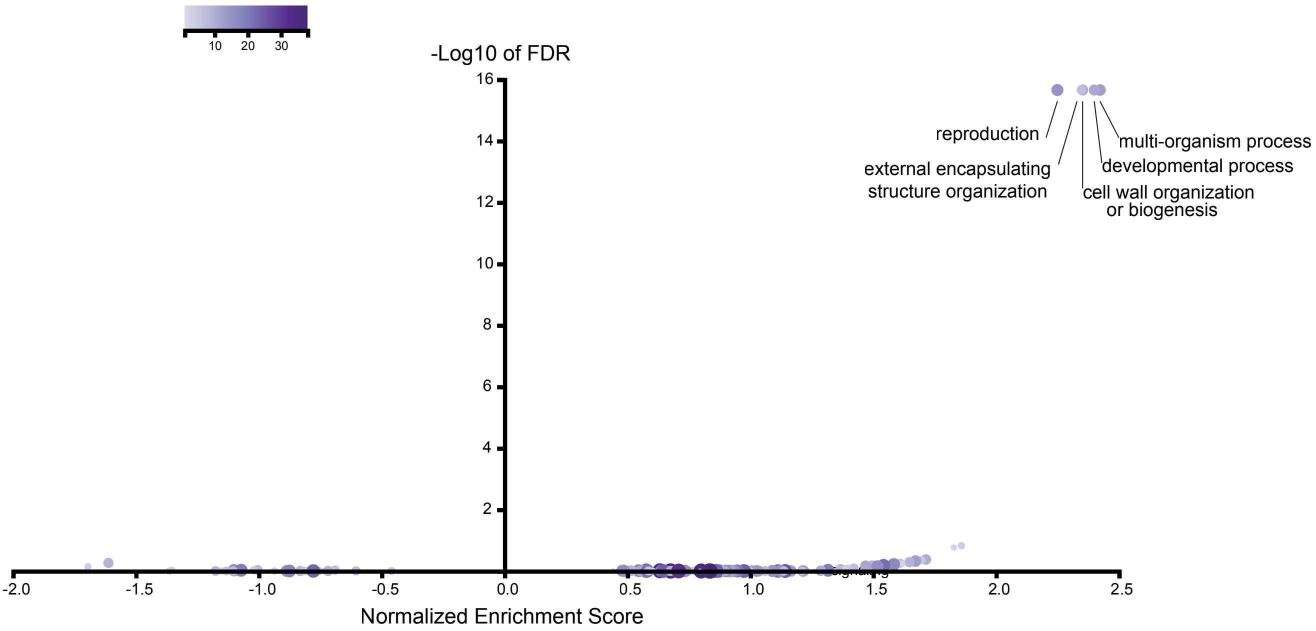
10
20
30
-Log10 of FDR
16
14
12
10
8
6
4
2
reproduction
external encapsulating
structure organization
multi-organism process
developmental process
cell wall organization
or biogenesis
-2.0
-1.5
-1.0
-0.5
0.0
0.5
1.0
1.5
2.0
2.5
Normalized Enrichment Score

Figure S2

**A**

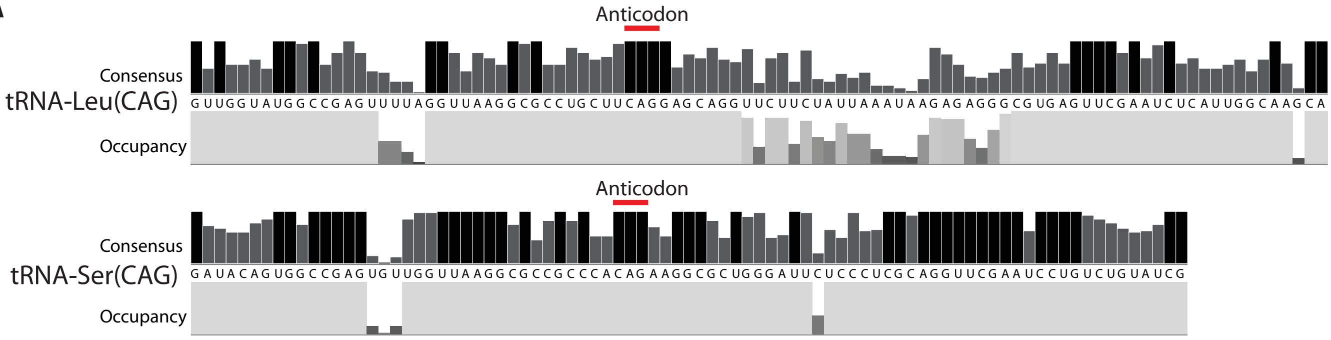


| | tRNA-Leu(CAG) | tRNA-Ser(CAG) |
|---|---|---|
| % columns 100% conserved: | 22.6% | 49.4% |

**B**

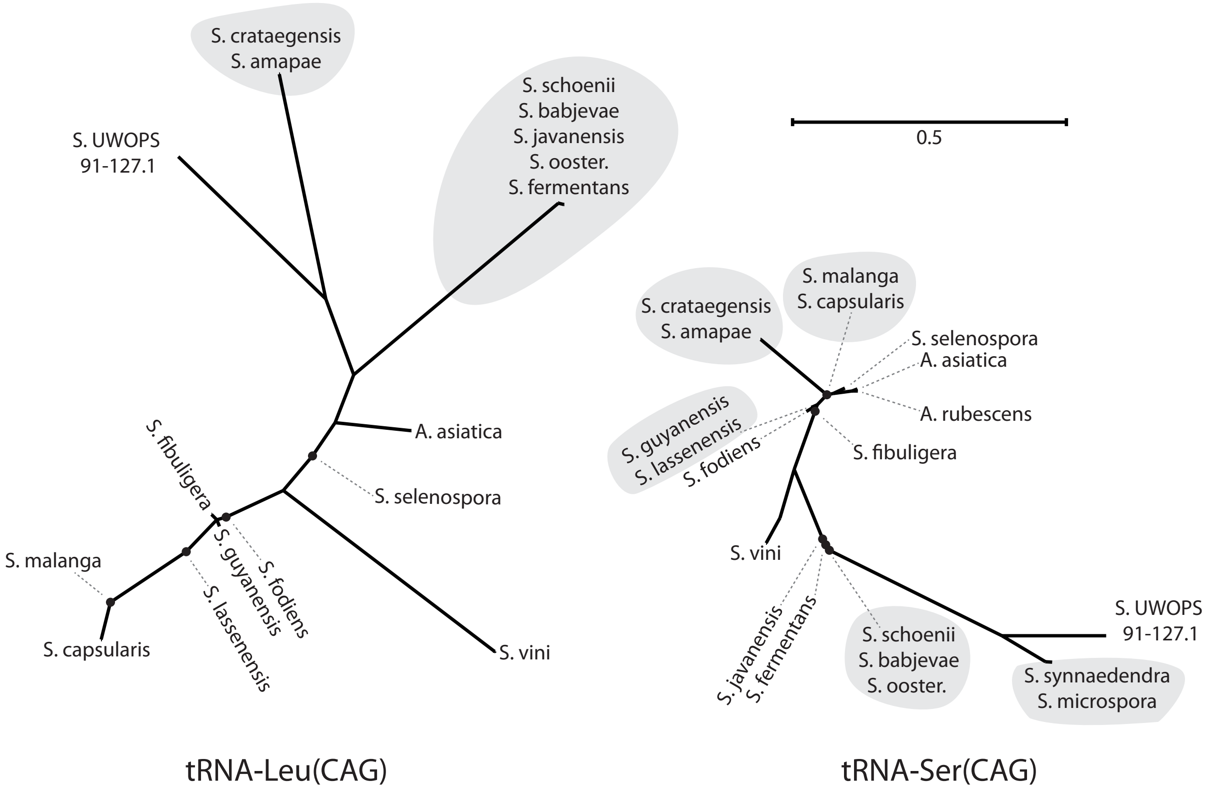


Figure S3

**Table S1.** Genome sequence data analyzed in this study.

| **Species** | **Strain** | **Assembly size (Mb)** | **Coverage** | **N50 (kb)** | **Reference** | **Accession number** |
|---|---|---|---|---|---|---|
| *S. capsularis* | NRRL Y-17639 (NT) | 17.8 | – | – | (Shen et al. 2018) | PPIG00000000 |
| *S. capsularis* | CBS 5064 | 17.3 | 40x | 283 | This study | PRJNA977123 |
| *S. capsularis* | CBS 7262 | 17.3 | 41x | 578 | This study | PRJNA977123 |
| *S. capsularis* | CBS 5638 | 17.0 | 35x | 562 | This study | PRJNA977123 |
| *S. malanga* | KCN 26 | 16.9 | – | – | Lee, unpublished. | CP025321 |
| *S. malanga* | JCM 7620 (T) | 16.7 | – | – | (Shen et al. 2018) | BCGJ01000000 |
| *S. fibuligera* | CBS 329.83 (T) | 22.3 | 32x | 5 | This study | PRJNA977123 |
| *S. fibuligera* | ATCC 36309 | 19.6 | – | – | (Choo et al. 2016) | CP015978 |
| *S. fibuligera* | MBY1320 | 19.1 | – | – | (Park and Kim 2021) | WXEZ01000000 |
| *S. fibuligera* | KPH12 | 19.6 | – | – | (Choo et al. 2016) | CP012823 |
| *S. crataegensis* | CBS 6447 (T) | 15.0 | 36x | 106 | This study | PRJNA977123 |
| *S. crataegensis* | CBS 6448 | 15.0 | 49x | 115 | This study | PRJNA977123 |
| *S. amapae* | CBS 7872 (T) | 15.0 | 52x | 132 | This study | PRJNA977123 |
| *S. vini* | CBS 4110 (T) | 17.4 | 37x | 30 | This study | PRJNA977123 |
| *S. vini* | CBS 4097 | 17.4 | 18x | 25 | This study | PRJNA977123 |
| *S. vini* | CBS 8420 | 17.4 | 32x | 25 | This study | PRJNA977123 |
| *Saccharomycopsis sp.* | UWOPS 91-127.1 | 12.2 | – | – | Wendland & Hesselbart, unpublished. | JNNM01000000 |
| *S. synnaedendra* | CBS 6161 (T) | 14.2 | 65x | 630 | This study | PRJNA977123 |
| *S. synnaedendra* | CBS 7763 | 15.9 | 57x | 1003 | This study | PRJNA977123 |
| *S. microspora* | CBS 6393 (T) | 15.8 | 58x | 524 | This study | PRJNA977123 |
| *S. schoenii* | CBS 7425 | 14.3 | – | – | (Junker et al. 2019) | JNFU00000000 |
| *S. schoenii* | CBS 7223 (T) | 13.5 | 61x | 161 | This study | PRJNA977123 |
| *S. oosterbeekiorum* | CBS 14943 (T) | 13.6 | 64x | 169 | This study | PRJNA977123 |
| *S. javanensis* | CBS 2555 (T) | 14.3 | 57x | 254 | This study | PRJNA977123 |
| *S. fermentans* | CBS 7830 (T) | 14.2 | 61x | 319 | This study | PRJNA977123 |
| *S. babjevae* | CBS 9167 (T) | 14.5 | 56x | 201 | This study | PRJNA977123 |
| *S. lassanensis* | CBS 8524 (T) | 12.9 | 68x | 320 | This study | PRJNA977123 |

| | | | | | | |
|---|---|---|---|---|---|---|
| *S. guyanensis* | CBS 12914 (T) | 13.9 | 65x | 792 | This study | PRJNA977123 |
| *S. fodiens* | CBS 8332 (T) | 14.9 | 59x | 770 | This study | PRJNA977123 |
| *S. selenospora* | CBS 2562 (T) | 14.8 | 56x | 136 | This study | PRJNA977123 |
| *A. asiatica* | JCM 7603 (T) | 20.3 | – | – | (Shen et al. 2018) | BCKQ00000000 |
| *A. asiatica* | CBS 457.69 (T) | 19.4 | 32x | 114 | This study | PRJNA977123 |
| *A. rubescens* | DSM 1968 (T) | 17.5 | – | – | (Riley et al. 2016) | LYBR01000000 |

In strain names, T denotes the type strain of a species, and NT denotes a neotype strain. Coverage and N50 data are shown for the genomes sequenced in this study.

**Table S2**. Sequences of oligonucleotides used in this study.

| Name | Primer sequence | Function |
|---|---|---|
| EOC1 | AATTCAACGCGTCTGTGAGG | screen upstream *ade2*::KanMX integration junction |
| EOC3 | ACGCTTACATTCACGCCCTC | screen downstream *ade2*::KanMX integration junction |
| EOC7 | aagttgttggcattctcggc | screen upstream *ade2*::KanMX integration junction |
| EOC8 | gcacaaacttctcgccgtat | screen downstream *ade2*::KanMX integration junction |
| EOC17 | cgagaagtctgacgtgcttacgattgagattgagcacgttaacgccgacaGCGACATGGAGGCCCAGAAT | generate RT *ade2*::KanMX |
| EOC18 | ggaagggtatatcttcaacgctgggttgttttgctccaagttgctcaagaCTTCGAGCGTCCCAAAACCT | generate RT *ade2*::KanMX |
| EOC29 | tacttagaaacatgcgtgct | tRNA-Leu locus PCR |
| EOC30 | gcaaggcacgtatagaaaac | tRNA-Leu locus PCR |
| EOC41 | aataggcgccatcgcgcatgtcaagacagcataaagacagcaaagcgcatGCGACATGGAGGCCCAGAAT | generate repair template tRNA-Leu::KanMX |
| EOC42 | caccagacgattcttactgagagtgtttatgcaaatatttcggcttgatgCTTCGAGCGTCCCAAAACCT | generate repair template tRNA-Leu::KanMX |

### References


Choo JH, Hong CP, Lim JY, Seo JA, Kim YS, Lee DW, Park SG, Lee GW, Carroll E, Lee YW, et al. 2016. Whole-genome de novo sequencing, combined with RNA-Seq analysis, reveals unique genome and physiological features of the amylolytic yeast *Saccharomycopsis fibuligera* and its interspecies hybrid. Biotechnol Biofuels 9:246.

Junker K, Chailyan A, Hesselbart A, Forster J, Wendland J. 2019. Multi-omics characterization of the necrotrophic mycoparasite Saccharomycopsis schoenii. PLoS Pathog 15:e1007692.

Park EH, Kim MD. 2021. Draft genome sequence data of the starch-utilizing yeast Saccharomycopsis fibuligera MBY1320 isolated from Nuruk. Data Brief 35:106888.

Riley R, Haridas S, Wolfe KH, Lopes MR, Hittinger CT, Goker M, Salamov AA, Wisecaver JH, Long TM, Calvey CH, et al. 2016. Comparative genomics of biotechnologically important yeasts. Proc Natl Acad Sci U S A 113:9882-9887.

Shen XX, Opulente DA, Kominek J, Zhou X, Steenwyk JL, Buh KV, Haase MAB, Wisecaver JH, Wang M, Doering DT, et al. 2018. Tempo and mode of genome evolution in the budding yeast subphylum. Cell 175:1533-1545 e1520.

Waterhouse AM, Procter JB, Martin DM, Clamp M, Barton GJ. 2009. Jalview Version 2--a multiple sequence alignment editor and analysis workbench. Bioinformatics 25:1189-1191.
